# Statistical inference of the Tree of Blobs of a phylogenetic network from quartet concordance factors

**DOI:** 10.64898/2026.05.28.728501

**Authors:** John A. Rhodes, Elizabeth S. Allman, Cecile Ané, Hector Baños

## Abstract

A phylogenetic network represents evolutionary relationships involving hybridization, gene flow, or admixture. While the full network may not be identifiable from genomic data under common coalescent models, its tree of blobs, depicting only the tree-like portions of the network structure, is. We introduce `ECToBlob` (Edge Contraction for Tree of Blobs), a new statistically-consistent algorithm to estimate the tree of blobs from quartet concordance factors. Starting from a resolved tree, `ECToBlob` successively contracts edges which statistical tests indicate do not belong in the tree of blobs, due to reticulate or polytomous signal. We show that `ASTRAL` provides a valid starting tree under common assumptions, in that, asymptotically in the number of loci, trees optimizing `ASTRAL`’s criterion refine the tree of blobs. We describe several algorithm variants, differing in how evidence from multiple tests are combined to determine if the edge should be contracted, and provide software implementations.

**Relevance to Life Sciences:** Hybridization, gene flow, or admixture are now recognized as important aspects of evolutionary history, but their genomic signal is confounded with that from a coalescent process, creating substantial challenges for inferring phylogenetic networks. The network’s tree of blobs identifies areas where reticulation occurred, separated by tree-like branching. `ECToBlob` quickly estimates the tree of blobs using quartet concordance factors from gene trees, and provides a measure of statistical support for its result. Performance is illustrated through simulation and on empirical data, using an implementation in the R package `MSCquartets`. While the presence of a blob may be all that can be inferred in some cases, in others `ECToBlob` offers a robust and principled way to focus further analyses on more local reticulate structure.

**Mathematical Content:** This work makes contributions to mathematical phylogenetics in optimization, combinatorics, and statistics. We show that any tree maximizing quartet support (the criterion underlying ASTRAL) is a refinement of the network’s tree of blobs under the coalescent model. Second, we give a concise proof that whether a network has a cut-edge corresponding to a given split is determined by information in certain subcollections of its 4-taxon subnetworks (quarnets). Finally, we propose valid statistical approaches for combining *p*-values across multiple quarnet hypothesis tests, proving that their use with specific decreasing test levels leads to statistically consistent inference as the number of loci grows.

**MSC codes:** 05C90, 60J95, 62-04, 62F07, 92D15

## 1. Introduction

This work offers a new statistically consistent algorithm to infer the tree of blobs of a species network from gene quartet data. The method requires as input a resolution of the tree of blobs which we show can be readily produced by available methods. Starting from this tree and using *p*-values of quartet hypothesis tests, it successively contracts edges that the statistical evidence suggests are not cut-edges of the network. It then returns an overall *p*-value for judging whether the remaining edges represent true cut-edges of the network. See Box 1.

### Box 1

**Overview**

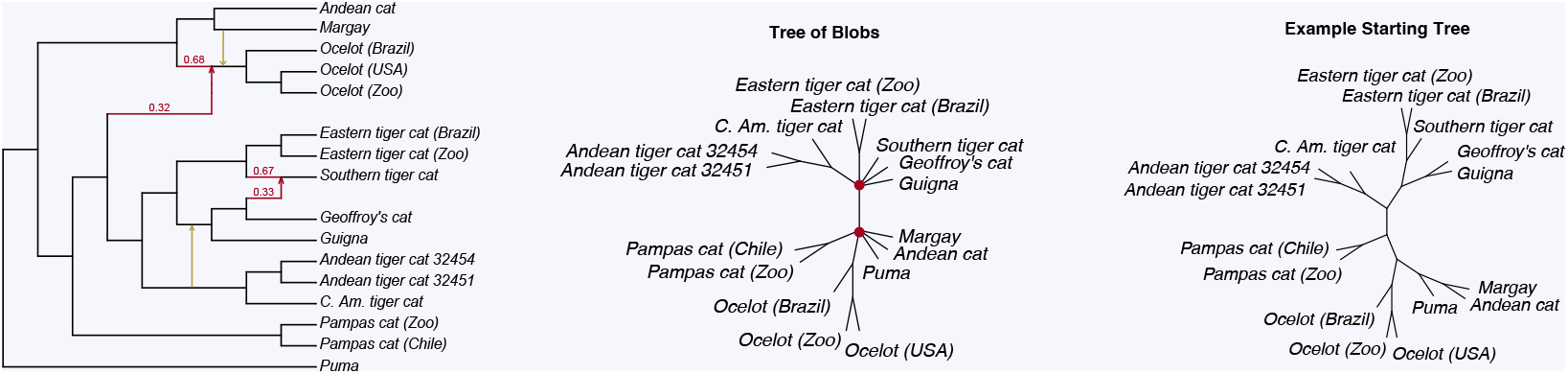

A phylogenetic network of Neotropical cats in the *Leopardus* genus [25] (left), showing putative evolutionary relationships that involve both tree-like descent and hybridization or admixture. Absent the gold reticulation edges, this network was inferred under a level-1 hypothesis [25], though subsequent analysis [7] suggests this might be too restrictive. Complex networks may not be inferrable from data, because some of the structure is theoretically nonidentifiable from particular data types. Inferring a network’s tree of blobs (center), which contracts the reticulate parts of the network to nodes, is a more feasible goal since identifiability is more readily established, requiring only the determination of cut edges in the true network. One approach is to infer a tree of blobs from a more resolved tree by testing individual edges, and contracting those that are unlikely to be cut edges. Any statistically valid resolution of the tree of blobs, such as the ASTRAL tree (right), can be used as an input starting tree for ECToBlob’s contraction procedure developed in this work.

The tree of blobs, *T*_*B*_(*N*), of a network *N* is the tree formed by contracting to nodes each multiconnected component of the network, that is, its non-tree-like substructures caused by reticulations. It has previously been shown to be identifiable from several types of data [2, 41, 33, 15], and the TINNiK algorithm (implemented in the `MSCquartetsR` package) [5, 32] provided the first statistically-informed approach to its inference. The tree of blobs isolates where reticulations have complicated the evolutionary history of species, and in some cases, may be the most detailed information we can reliably extract from datasets. In other cases, it may lead to subsequent analyses focused on determining each blob’s finer structure. Indeed, the NANUQ+ algorithm pioneered this approach for level-1 network inference [7].

The tree of blobs inference method developed here begins with a tree *T*_0_ which is a resolution of the tree of blobs *T*_*B*_(*N*). We prove that the widely-used species tree inference method `ASTRAL` [43], or any method with the same optimality criterion, provides such a tree (up to heuristic elements of its search method) under several generative gene tree models, including the widely adopted Network Multispecies Coalescent (NMSC) model with independent inheritance. Note that applying any species tree inference method to data from a network is, in general, a case of model misspecification, as there is no “species tree” to infer and *a priori* it is unclear what might be produced. In fact, networks are known where the optimality criterion of `ASTRAL` produces trees converging to ones *not* displayed on the networks as the amount of data increases [14, 37]. By proving that `ASTRAL`’s criterion leads to a resolution of the tree of blobs, we show that, despite the fact that its output should not be called “the species tree,” such a method can still be the first step for consistent inference of *T*_*B*_(*N*), and hence of *N*.

With a resolution of the tree of blobs in hand, our method then uses hypothesis tests for hybridization or admixture at the quartet (4-taxon) level, as developed in [30]. While other tests could be used in their place, both theory and the simulation study of [11] suggest that these tests gives superior performance in a wide range of circumstances. With a null hypothesis of the quartet topology displayed on *T*_0_, we obtain *p*-values which, if small, suggest that certain edges are not present in the tree of blobs. Since we perform many such tests, which are not independent and have unknown covariance, *p*-values must be combined with care due to multiple testing issues. We propose and test combining *p*-values using the well-established Bonferroni-adjusted minimum approach, the Cauchy combination test of [26], and its variants introduced by [12]. While Cauchy combination tests have previously been used in phylogenomics to determine whether tree or network inference is more appropriate for a dataset [20], our use is novel. We successively contract edges of *T*_0_ with the smallest combined *p*-values to estimate *T*_*B*_(*N*), stopping when a combined overall *p*-value for the multifurcating tree rises above a level *α*.

In addition to the choice of multiple test correction methods, we propose three ways of choosing 4-taxon sets to test each edge in the contracting process from *T*_0_. We prove that all these variants of the algorithm are statistically consistent under mild conditions, provide software implementation in the R package `MSCquartets`, and demonstrate performance on simulated and empirical data.

When this work was well underway, we learned a different group was independently pursuing the same outline for tree of blobs inference, implemented in `TOB-QMC` [13]. Our work differs by emphasizing a more statistical viewpoint in the treatment of hypothesis test results, as well as strengthening mathematical aspects of the formal justification. We discuss these differences and more in the concluding section of this work.

Section 2 recalls definitions and terminology. Section 3 proves that under common gene tree distribution models on a species network, including the NMSC model with independent inheritance, maximizing gene quartet concordance with proposed trees selects a resolution of the tree of blobs. In Section 4, we establish a fundamental combinatorial result on the presence of a cut edge in a network, based on cut edges in a subset of 4-taxon subnetworks. This result can be used to verify whether a possible split does not appear in a network’s tree of blobs. Hypothesis testing for this quartet property is the subject of Section 5. Then in Section 6, we introduce several statistically justified means of combining *p*-values from non-independent tests, establishing some of their basic properties. Our algorithm for tree of blobs inference appears in Section 7, while Section 8 establishes its statistical consistency. After a demonstration of performance on both simulated and empirical data in Section 9, we conclude with a discussion in Section 10.

## 2. Preliminaries

We adopt standard definitions of metric rooted phylogenetic networks *N* ^+^ and semidirected phylogenetic networks *N*^−^ found in [3], using *N* to denote either. Specifically, we allow for the possibility of non-binary internal nodes, and for parallel edges, as such networks arise when considering the subgraph induced by a subset of taxa. An internal tree node has 1 incoming edge, and possibly 1, 2, or more outgoing ones. A hybrid node has 2 or more incoming edges, and 1 or more outgoing ones.

Unless otherwise noted within arguments, we assume all our networks are LSA networks, meaning the root coincides with the least stable ancestor (LSA) of the full set of taxa. By removing edges and nodes ancestral to the LSA, any network can be made to have this property. Since methods that utilize only topological gene trees to infer species network structure under standard models, as this work does, can provide no information about network features above the LSA, there is no loss in reducing to LSA networks.

Given a rooted network *N* ^+^ on taxa *X*, by the *induced subnetwork* (or *induced subtree*) 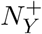 on a subset of taxa *Y* ⊂ *X* we mean the network obtained by retaining only those edges ancestral to taxa in *Y* below the LSA of *Y*, and suppressing degree-2 nodes. If |*Y* | = 4, 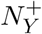 is called a *quarnet*. The induced subnetwork differs from the induced subgraph on *Y* which retains all edges ancestral to *Y*, neither omitting those above *Y* ‘s LSA nor suppressing degree-2 nodes. Induced semidirected networks (including unrooted trees) can be obtained from induced rooted ones, but can also be defined directly [41].

### Blobs

A *blob* in a phylogenetic network *N* is a maximal 2-edge connected subgraph, and a *trivial* blob is a blob that contains a single node. When a network is binary, a non-trivial blob is equivalently a maximal biconnected (i.e., 2-node connected) subgraph. Note that in a non-binary network with two biconnected components sharing a single node, these components are in the same blob by our definition (see [33, Figure 1]). An *n-blob* is one incident to *n* cut edges of the network.

**Figure 1.**
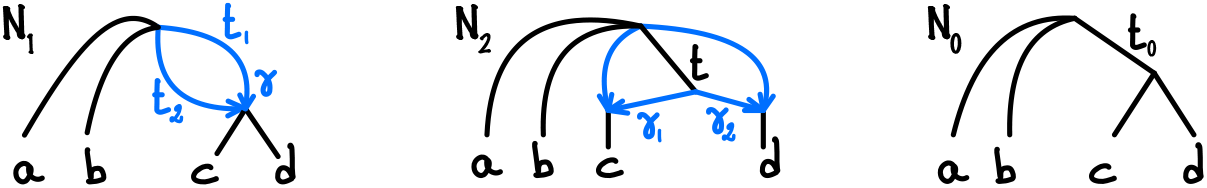
*Under the NMSC models, these networks satisfy p*_*ab*|*cd*_ > *p*_*ac*|*bd*_ = *p*_*ad*|*bc*_, *introduced later as* (5.2) *in model Assumption 1. N*_1_ *(left) and N*_2_ *(middle) are not binary and do not have a non-trivial cut-edge split. Only N*_0_ *(right) does. If t*_1_ > 0 *or t*_2_ > 0 *then N*_1_ *satisfies* (5.2) *and has the same quartet CFs as N*_0_ *with an appropriate t*_0_ > 0 *(see [8, Example 1]). N*_2_ *satisfies* (5.2) *with the same quartet CFs as N*_0_ *with t*_0_ *satisfying* (1 − *e*^−*t*0^) = *γ*_1_*γ*_2_(1 − *e*^−*t*^).

### The tree of blobs

By the *tree of blobs T*_*B*_(*N*) of a network *N*, we mean the (unrooted) tree obtained by contracting each blob of *N*^−^ to a node, and then *reducing* by suppressing degree-2 nodes. For a rooted network, the tree of blobs is obtained by first passing to its semidirected form (suppressing the root if it has degree 2) and then to the tree of blobs. The tree of blobs of a network typically has multifurcations (polytomies) arising from the network’s *m*-blobs for *m* ≥ 4. However, 3-blobs in a network *N* lead to degree-3 nodes in *T*_*B*_(*N*), and these nodes of *T*_*B*_(*N*) are indistinguishable from those nodes arising from binary nodes in *N*. A network’s 2-blobs, however, are simply lost as the degree-2 node to which they are contracted is suppressed.

By a *resolution* of a tree *T*, we mean any tree 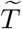 from which *T* can be obtained via edge contractions. A resolution therefore, may or may not be binary, and in particular, a resolution of a tree of blobs may still have some multifurcations, including ones that only partially reflect blobs of *N*.

### Quartets

By a *quartet* for taxa *a, b, c, d* we mean any of the unrooted resolved topological trees *ab*|*cd, ac*|*bd, ad*|*bc* or the unresolved star tree denoted *abcd*. A quartet is *displayed* on a tree *T* on taxa *X* if it is the unrooted tree on the 4 taxa induced from *T*. Under many gene tree distribution models (e.g., the NMSC), gene tree quartets must be resolved, though a 4-taxon species network may be unresolved or have a reticulate topology.

A quartet *ab*|*cd* displayed on *T defines* an edge *e* of *T* if, when *e* and its incident edges are removed from *T, a, b, c, d* are in distinct connected components of the resulting graph. We refer to the partition of *X* according to these connected components as the *multipartition* induced by *e*, or if *T* is binary as the *quadripartition*.

Given a multiset 𝒬 of quartet trees on a set of taxa *X*, the unweighted *Maximum Quartet Support (MQS) criterion* is an optimality criterion for choosing a tree (or trees) *T* on *X*, by maximizing the number of elements of 𝒬 displayed on *T*. The software `ASTRAL` [29], for instance, uses this criterion and a heuristic search over tree space, working from a collection of gene trees which are (conceptually, though not in practice) summarized by their displayed quartets. `TREE-QMC` [19], though algorithmically different, uses the same criterion and data. Both provide fast and statistically consistent species tree inference under various models (but in the absence of gene flow modeled by networks) [28, 21, 37, 19]. Variants of MQS allow for weighting of quartets, with the weight perhaps measuring support or confidence.

### Models and concordance factors

Various models have been proposed which assign probabilities to gene trees (or more properly, non-recombining locus trees) arising on a species network *N* ^+^. The (independent) Network Multispecies Coalescent (NMSC) is built on the Kingman coalescent, constrained to lineages within network edges, with inheritance probabilities determining independent lineage behavior at reticulations. The NMSC with common inheritance constrains all lineages for a gene to follow the same inheritance behavior at reticulations, while the correlated inheritance NMSC interpolates between these two models. Under the Displayed Tree model the realizable gene trees are those displayed on the network, with probabilities determined by the relevant hybridization parameters. For more details on these models, see e.g. [33, Def. 12] or [3, Def. 8]. Note that all of these models treat individual gene trees as independent, and behave well under marginalization over taxa (equivalently, under passing to subnetworks). We will not limit our arguments to any one of these models, instead referring to key properties of the distributions that we require, and pointing out which common models have them.

A *quartet Concordance Factor* (CF) associated to a gene tree distribution is simply the vector of probabilities of the resolved gene tree topologies for a set of 4 taxa:

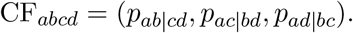

For standard coalescent models, the entries of this CF add to 1, as the probability of an unresolved gene tree is 0. For the displayed tree model this need not be the case, since an induced 4-taxon network might be a star tree.

Because gene trees are not observable, in practice we use inferred gene trees as ‘data.’ Assuming a correctly specified model, as sequence length grows standard methods of inference from sequences will, with probability approaching 1, return the correct topology. Thus CFs can be estimated by the empirical frequencies of quartets across a sample of gene trees. For the displayed tree model, it is convenient to redefine the CF in the star case as (1*/*3, 1*/*3, 1*/*3), and treat unresolved inferred trees as contributing this to quartet topology counts. More formally, define the *empirical quartet Concordance Factor* 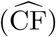 as the frequencies of the resolved quartet trees displayed on a collection of gene trees, where a star tree contributes (1*/*3, 1*/*3, 1*/*3). Then, for all our models, 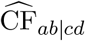 converges to CF_*ab*|*cd*_ as the number of loci approaches infinity for a true sample of gene trees from the model. It also holds for gene trees inferred from sequences of growing length, provided that the gene tree inference method for a true polytomous quartet either returns the correct star topology or is unbiased in which resolution it returns.

Hypothesis testing of whether empirical CFs are in accord with theoretical ones can play a valuable role in network inference. For 4 taxa, the *quartet count Concordance factor* (qcCF) is the vector of counts of the quartet topologies across a set of gene trees. This is the data summary on which such hypothesis tests are performed.

However, it is known that under the models mentioned above, the presence of a biconnected component with one entry node and one exit node is never identifiable from quartet CFs in the very strong sense that replacing the component by a single edge of an appropriate length, and making no other changes to the network, will leave all quartet CFs unchanged [36, 10, 8]. For example, the network *N*_1_ of Figure 1 (left) is indistinguishable by CFs from *N*_0_ (right) in which a cut edge replaces the parallel edges, despite the trees of blobs of *N*_1_ and *N*_0_ being unresolved and resolved, respectively.

Since our algorithm is based on quartet CFs, this point has implications for exactly what can be inferred as a tree of blobs. If a network has a 2-blob with one in-edge and one out-edge, then replacing that blob with an edge has the same topological impact on the tree of blobs as contracting it to a node, as degree-2 nodes are suppressed. However, if a biconnected component has exactly one entry node and one exit node, and if both are adjacent to 2 or more cut-edges, then replacing the component by an edge may increase the resolution of the tree of blobs, as discussed above. We return to this point later, and impose a technical assumption of no multifurcations within blobs to address it.

## 3. Resolving the tree of blobs

As our method requires a resolution *T*_0_ of a network’s tree of blobs as input, we first show that established methods can produce one. Specifically, we show that, with mild assumptions on the gene tree distribution model, any resolved tree that maximizes the sum of probabilities of displayed quartets on gene tree must be a resolution of *T*_*B*_(*N*). A straightforward probability argument then implies that any algorithm that seeks from a collection of gene trees a tree *T* satisfying the MQS criterion yields, with probability going to 1 as the number of gene trees increases, a resolution of *T*_*B*_(*N*).

While the MQS criterion is often used to infer a species tree from gene trees, we reiterate that its use here, on data generated from a network with reticulations, should not be described as “inferring the species tree.” In fact, it is known that under the NMSC model and ideal conditions an optimal MQS tree need not even be displayed on the network [14] for a positive measure subset of the parameter space.

While we do not attempt to address how MQS resolves a network’s blobs, we make several simple observations. Under the NMSC and other models, one can extend the model parameterization to a continuous function allowing for *γ*s in the interval [0, 1]. Since *γ*s of 0 and 1 correspond to displayed trees, continuity implies that for *γ*s sufficiently close to 0 and 1, MQS will infer displayed trees as well. Thus, for a positive-measure subset of parameter space MQS will select a displayed tree of the network. Varying *γ*s continuously between choices of parameters leading to different trees also shows that there must be parameter values for which multiple optimal MQS trees exist.

Note that while Habib et al. [18] considered when MQS and similar criteria lead to terraces of trees with tied scores, those results focus on gene trees with missing taxa, application of the optimization criterion to datasets of fixed finite size, and gene trees not produced from a particular model. We consider none of these, working only in the ideal situation of no missing taxa and arbitrarily large datasets of gene trees.

The main goal of this section is to show the following.

### Theorem 3.1.

*Let 𝓊* (*X*) *denote the set of unrooted topological phylogenetic trees on a taxon set X, and N* ^+^ *a rooted metric phylogenetic network on X. Consider any gene tree distribution satisfying:*

1. *If there exists a cut edge of N* ^+^ *separating taxa a, b from c, d, then* (3.1)
2. *For any x, y, z, w* ∈*X, p*_*xy*|*zw*_ *is determined by the network on x, y, z, w induced from N* ^+^, *but is independent of both the metric and topological structure of pendant chains of 2-blobs and edges leading to single taxa*.

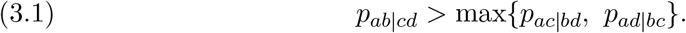

*Then any T* ∈ 𝓊 (*X*) *maximizing the sum over quartet trees displayed on T of the probabilities of gene tree quartets is a resolution of the tree of blobs of N* ^+^. *That is, if T*_*B*_(*N* ^+^) *is the (reduced unrooted) tree of blobs of N* ^+^, *and*

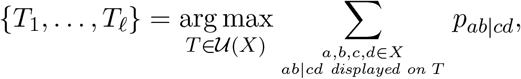

*then every split associated to an edge of T*_*B*_(*N* ^+^) *is associated to an edge of every T*_*i*_.

### *Remark* 3.2.

For a model in which only fully resolved gene trees have positive probability, such as coalescent models, all maximizers *T*_*j*_ in Theorem 3.1 must be fully resolved, even if *N* ^+^ is not binary, since any multifurcation in a potential maximizer can be arbitrarily resolved to increase the MQS score. For the displayed tree model, this is not the case, as resolving a polytomy on *T* arising from one in the network will have no impact on *T* ‘s MQS score. If, as suggested in Section 2, we treat each resolution of a polytomy as having equal probability 1/3 (as is appropriate if gene trees are inferred), then maximizers will be binary.

### *Remark* 3.3.

The two assumptions of the theorem are met by common models in phylogenomics. The NMSC model with independent or correlated inheritance satisfies (3.1) provided the network has no anomalous quartets and that cut edges (which are necessarily tree edges) have positive lengths [8]. The NMSC with common inheritance, and the displayed tree model (with no coalescent process) also satisfy (3.1) provided only that cut edges have positive lengths. All of these models also satisfy assumption 2 of Theorem 3.1.

### *Remark* 3.4.

Under models with gene duplication and loss, (3.1) is also known to hold when the species phylogeny is a tree, either when a single gene copy is sampled per individual, or when all gene copies from each individual are treated as multiple alleles [24, without ILS], [27, 21, with ILS]. However, it remains an open question whether these results hold for duplication and loss models when the species phylogeny has reticulations.

Our argument uses the following.

### Lemma 3.5.

*Let* 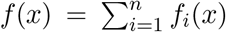 *be a real-valued objective function to be maximized over a set U. Suppose there exists an x*_0_ ∈ *U which simultaneously maximizes each of the f*_*i*_. *Then x*_0_ *maximizes f, and any other maximizer of f also maximizes each of the f*_*i*_.

*Proof*. Since each of the *f*_*i*_ takes on no values larger than *f*_*i*_(*x*_0_), it is immediate that *x*_0_ maximizes *f*. If *x*_1_ is another maximizer of *f*, yet *f*_*i*_(*x*_1_) < *f*_*i*_(*x*_0_) for some *i*, then 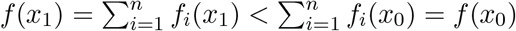, which is a contradiction.

We now establish the theorem above.

### *Proof of Theorem 3.1*.

For any 4-taxon set *Q* ⊂ *X*, the induced subtree on *Q* of *T*_*B*_(*N* ^+^) is either a resolved tree or a star tree. Let ℛ be the collection of *Q* for which the tree is resolved, and 𝒮 the collection for which it is a star. Then 𝒮 can be further decomposed as 𝒮 = ⊔_*m∈ℳ*_ 𝒮_*m*_ where ℳ is the set of multifurcations (nodes of degree *k* ≥ 4) on *T*_*B*_(*N* ^+^), and 𝒮_*m*_ is the collection of those *Q* whose induced subtree has *m* as its central node. Then the objective function

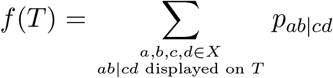

can be expressed as

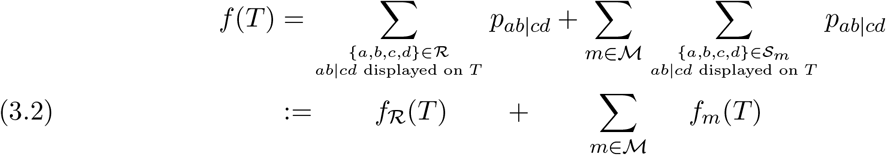

We next investigate maximizers of *f*_*R*_ and of *f*_*m*_ separately, in order to apply Lemma 3.5.

Consider first *f*_ℛ_. If {*a, b, c, d*} ∈ ℛ and *ab*|*cd* is the quartet displayed on *T*_*B*_(*N* ^+^) then, by assumption (3.1), *p*_*ab*|*cd*_ > max{*p*_*ac*|*bd*_, *p*_*ad*|*bc*_}. Thus *T*_*B*_(*N* ^+^) is a maximizer of *f*_ℛ_ and any maximizer of *f*_ℛ_ must display the same resolved quartets as does *T*_*B*_(*N* ^+^). These maximizers are thus all resolutions of *T*_*B*_(*N* ^+^).

Now consider *f*_*m*_ for a fixed multifurcation *m* ∈ ℳ. Let *X*_1_, *X*_2_,…, *X*_*k*_ be the partition of *X* according to the connected components of the graph obtained by deleting *m* from *T*_*B*_(*N* ^+^). Then {*a, b, c, d*} ∈ *S*_*m*_ exactly when *a, b, c, d* are in distinct *X*_*i*_. Moreover, by assumption 2, *p*_*ab*|*cd*_ depends only on the *X*_*i*_ containing *a, b, c, d*, so we denote these probabilities by *pX*_*g*_ *X*_*h*_|*X*_*i*_*X*_*j*_. Thus

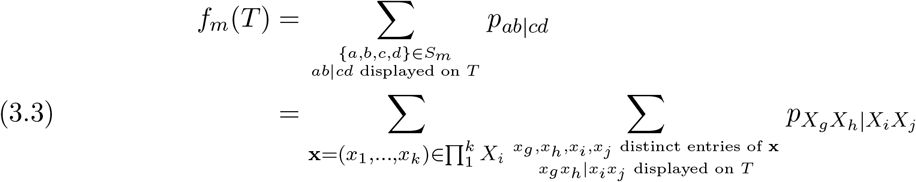

Viewing the *X*_*i*_ as taxon labels, for trees 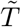 on 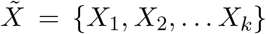 define the objective function 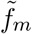 by

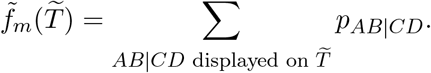

Suppose 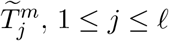, are all the maximizers of 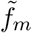. Taking any 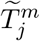 and attaching arbitrary rooted subtrees on the taxa in *X*_*i*_ at the leaf of 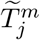 labelled by *X*_*i*_ gives a tree that simultaneously maximizes the inner sum of (3.3) for each **x**. Thus by Lemma 3.5, every maximizer *T*_*m*_ of *f*_*m*_ must be a tree that, when restricted to any choice of representatives of the *X*_*i*_, has one of the topologies 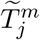. However, at this point, we do not know that different choices of representatives in the *X*_*i*_ all are associated to the same 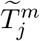.

For each *m* ∈ ℳ, pick a maximizer 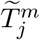 of 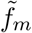 and ‘splice’ these into *T*_B_ (*N*^+^) as follows:

Remove *m* and for each edge *e* = {*m, x*_*i*_} incident to *m* leading to *X*_*i*_ in *T*_*B*_(*N* ^+^), remove *e* and identify *x*_*i*_ with the node *X*_*i*_ in 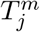. This gives a tree that maximizes all terms in (3.2), so by Lemma 3.5 it maximizes *f*. Moreover, any maximizer *T* of *f* must be obtained in this way. This is because maximizing *f*_ℛ_ requires that for each *m* ∈ ℳ a maximizer *T* must have cut edges separating each of *m*’s *X*_*i*_ from the other taxa. Therefore, for each *m*, any choice of representatives of the *X*_*i*_ must give the same induced subtree 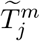.

### Corollary 3.6.

*Given n gene trees on a taxon set X produced under a model satisfying the assumptions of Theorem* 3.1, *the optimal tree(s) under the MQS criterion will, with probability* → 1 *as n* → ∞, *be resolutions of the tree of blobs of N* ^+^.

*Proof*. For each set of 4 taxa, the empirical quartet concordance factors, 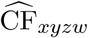 as defined in Section 2, converge in probability to the true CF_*xyzw*_ as *n* → ∞. Since there is a deterministic and finite number of choices of 4 taxa, this is uniform across 4-taxon sets, and for any *T*

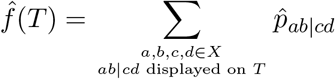

converges in probability to *f* (*T*), the objective function in Theorem 3.1. Since the set of candidate trees 𝓊 (*X*) is finite, the probability that 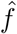 and *f* have different sets of optima → 0 as *n* → ∞.

Note that current software, such as `ASTRAL,` seeking to maximize the MQS criterion from a collection of gene trees returns only one tree, and not necessarily all optimal trees. In general, we cannot say that any algorithm giving a single MQS optimal tree must, on growing datasets, give a sequence of trees converging to a resolution of the tree of blobs, since such a sequence could have several accumulation points (trees).

## 4. Combinatorics of quartets and cut-edge splits

The key combinatorial understanding that justifies the algorithms we develop to infer the tree of blobs *T*_*B*_(*N*) concerns the relationship between quartet networks induced by *N* and the splits associated to the cut edges of *N*. While it is clear that such a network split implies that many induced quarnets also have a cut edge, for our work we need a converse, which we prove below in Theorem 4.1. It will allow for evidence from the quarnets, as judged by hypothesis tests, to be used to determine which edges in a resolution of the tree of blobs should be contracted since they do not arise from a cut edge of the network.

To decide if an edge *e* in a resolution *T* of the tree of blobs of *N* ^+^ should be contracted, there are at least three natural collections of 4-taxon sets one might consider. One is those sets *defining the edge e*, in the sense that the internal edge of the quartet tree induced by the 4 taxa on *T* is exactly *e*. Since for a binary resolution this means the 4 taxa are drawn from the quadripartition of the taxa formed by deleting *e* and its endpoints, we call these the *quadripartition sets* of *e*, denoting the collection by 𝒮_quad_ (*e*). If either end of *e* is a multifurcation (i.e., has degree >3), then we can similarly consider all those 4-taxon sets defining the edge with taxa drawn from different sets in the multipartition of the taxa induced by deleting *e* and its endpoints, i.e, the collection of *multipartition sets*, 𝒮_mul_ (*e*). Finally we also consider the *bipartition sets*, forming the collection 𝒮_bi_ (*e*), in which 2 taxa are drawn from each block of the bipartition of taxa defined by deleting *e* but keeping its endpoints. These 4-taxon sets define paths in *T* that include *e*, but may include other edges as well. Note that 𝒮_quad_ (*e*) only makes sense if *e* has degree-3 endpoints, and in that case 𝒮_quad_ (*e*) = 𝒮_mul_ (*e*). We will later refer to the options of the `bi, mul`, or `quad` quartet sets as the choice of a *quartet type*.

In this section, we establish that if *e* in *T* should be contracted to obtain the network’s tree of blobs, then each of 𝒮_quad_ (*e*), 𝒮_mul_ (*e*), 𝒮_bi_ (*e*) must contain a 4-taxon set whose quartet induced from *T* conflicts with its quarnet induced from *N*^+^, in the sense that its quarnet either has no non-trivial *cut-edge split*, or has a non-trivial cut-edge split different from that in *T*. Here we call a split *A*|*B* a cut-edge split of *N* ^+^ if the network has a cut edge *e* whose removal disconnects the graph into connected components with taxon sets *A* and *B*. This contrasts with a related notion: A split *A*|*B* is *displayed* on *N* if *A*|*B* is a cut-edge split of some tree *T* displayed in *N*.

For a rooted network *N* ^+^ on *X* and taxon *a* ∈ *X*, the *funnel of a* is the connected subgraph whose edges are those of *N* ^+^ which are ancestral to *a* and to no other taxon. *N* ^+^ can be viewed as its induced subgraph on *X* ∖ {*a*} together with the funnel of *a attached* at their common nodes. (Recall that the induced subgraph retains degree-2 nodes, as well as any edges and nodes above the LSA of the taxa.)

The following general result links cut-edge splits in the full network to those in quarnets.

### Theorem 4.1.

*Let N* ^+^ *be a rooted network on X (not necessarily binary), and A*|*B a non-trivial split of X. Let A* = *A*_1_ ⊔ *A*_2_, *B* = *B*_1_ ⊔ *B*_2_ *be arbitrary partitions into non-empty sets. If for all a*_1_ ∈ *A*_1_, *a*_2_ ∈ *A*_2_, *b*_1_ ∈ *B*_1_, *b*_2_ ∈ *B*_2_, *a*_1_*a*_2_|*b*_1_*b*_2_ *is a cut-edge split of the induced 4-taxon network* 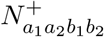, *then A*|*B is a cut-edge split of N* ^+^.

*Proof*. Under the hypothesis of the claim, choose *ã*_1_ ∈ *A*_1_, *ã*_2_ ∈ *A*_2_, 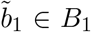, 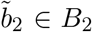 so that the rooted 4-taxon induced subgraph 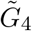 (without suppressing degree-2 nodes or removing anything above the LSA) on these taxa has the minimal number of cut edges inducing 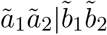 (see Figure 2). Then the induced subnetwork 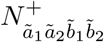 has two 3-blobs.

**Figure 2.**
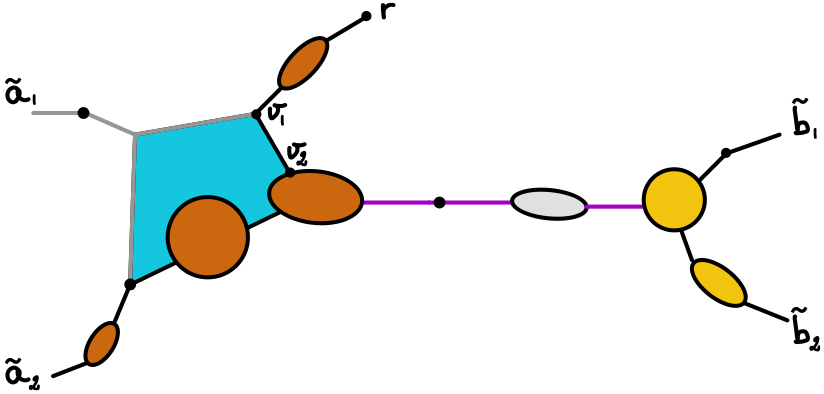
*Illustration for the proof of Theorem* 4.1, *showing a possible induced subgraph* 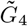 *on taxa ã*_1_, *ã*_2_, 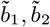, *including the root r of the full network, and bounded areas for non-trivial blobs. Straight black and purple edges are the cut edges in* 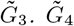 *is the union of* 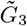 *and the funnel of ã*_1_, *which is shown with the blue area and grey edges. The LSAs of* {*ã*_1_, *ã*_2_, 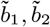} *and of* {*ã*_2_, 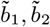} *are shown as v*_1_ *and v*_2_ *respectively. Purple edges are those in E, inducing* 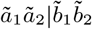. *The ã*_2_*-side of* 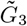 *is shown with brown blobs, and the* 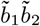 *-side with yellow blobs*.

Since 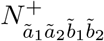 has a cut-edge split 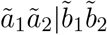, up-down paths from *ã*_1_ or *ã*_2_ to a 3-blob lead to one of these blobs first, while up-down paths from 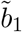 or 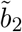 lead to the other. Moreover, the subgraphs between each taxon and its closest 3-blob have the form of a chain of 2-blobs and cut edges, as does the subgraph connecting the two 3-blobs.

Let *E* be the non-empty set of cut edges in 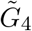 inducing the split 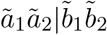. By choice of *ã*_*i*_, 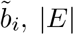, |*E*| is minimal. Let 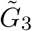 be the subgraph of *N* ^+^ (or of 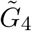) induced by *ã*_2_, 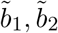. Since *G*_3_ also contains the edges in *E*, we define the *ã*_2_*-side* of that subgraph as the connected component in which *ã*_2_ lies when the edges in *E* are deleted, and similarly the 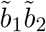 *-side*.

Viewing 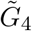 as the graph obtained from 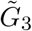 by attaching the funnel of *ã*_1_, and similarly, considering the subgraph induced by *a*_1_, *ã*_2_, 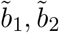, with *a*_1_ ∈ *A*_1_, *a*_1_≠ *ã*_1_ as 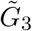 with the funnel of *a*_1_ attached, the minimality of |*E*| implies that the funnel of *a*_1_ can only attach to 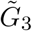 on the *ã*_2_-side. Since this is true for all *a*_1_ ∈ *A*_1_, the edges in *E* induce the split 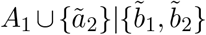 on the subnetwork on those taxa. Reasoning similarly with *ã*_2_, 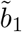, and 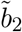 shows the edges in *E* induce the split *A*|*B* on *N* ^+^.

Taking *A*_1_ and *B*_1_ to be singleton sets immediately yields the following, relevant to bipartition sets.

### Corollary 4.2.

*Let N* ^+^ *be a network on X, and A*|*B a non-trivial split of X. Then either A*|*B is a cut-edge split of N* ^+^ *or for any choice of taxa a*_1_ ∈ *A and b*_1_ ∈ *B, there exist distinct taxa a*_2_ ∈ *A and b*_2_ ∈ *B such that a*_1_*a*_2_|*b*_1_*b*_2_ *is not a cut-edge split of the induced network* 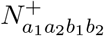 *on a*_1_, *a*_2_, *b*_1_, *b*_2_. *That is, either* 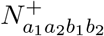 *has a 4-blob and thus no non-trivial cut-edge splits, or it has one of the cut-edge splits a*_1_*b*_1_|*a*_2_*b*_2_ *or a*_1_*b*_2_|*a*_2_*b*_1_.

In the case of binary networks, this corollary follows from Lemma 1 of [17], which depends on results in [22]. Our proof is considerably briefer and we relax the binary condition, extending part of Lemma A.1 of [33] to networks with more than 4 taxa.

For quadri- and multipartition sets of 4 taxa, we obtain the following.

### Corollary 4.3.

*Let N* ^+^ *be a network on X, and A*|*B a non-trivial split of X, with finer partitions* 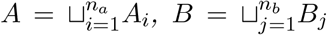 *into non-empty sets (n*_*a*_ ≥ 2, *n*_*b*_ ≥ 2*). Then either A*|*B is a cut-edge split of N* ^+^ *or for any choice of sets A*_*i*_ *and B*_*j*_, *there exist taxa a*_1_ ∈ *A*_*i*_, *a*_2_ ∈ *A* ∖ *A*_*i*_ *and b*_1_ ∈ *B*_*j*_, *b*_2_ ∈ *B* ∖ *B*_*j*_ *such that a*_1_*a*_2_|*b*_1_*b*_2_ *is not a cut-edge split in the induced network N*_*a*1_*a*_2_*b*_1_*b*_2_ *on a*_1_, *a*_2_, *b*_1_, *b*_2_. *That is, either N*_*a*1_*a*_2_*b*_1_*b*_2_ *has a 4-blob and thus no non-trivial cut-edge splits, or it has exactly one of the cut-edge splits a*_1_*b*_1_|*a*_2_*b*_2_ *or a*_1_*b*_2_|*a*_2_*b*_1_.

*Proof*. This follows from Theorem 4.1 by taking 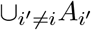 as *A*_2_ and 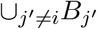 as *B*_2_ in that statement.

Unlike Corollary 4.2, Corollary 4.3 does not allow the specification of 2 specific taxa in blocks *A*_*i*_ and *B*_*j*_, only the two blocks themselves. That the conclusion need not hold if the taxa were specified is easily seen through a small example. Consider a 5-cycle sunlet network on taxa *a, b, c, d, e*, with *a* the hybrid descendant. Set *A*_1_ = {*a, b*}, *A*_2_ = {*c*}, *B*_1_ = {*d*}, *B*_2_ = {*e*}, and specify the taxa *b* in *A*_1_, and *d* in *B*_1_. Then no quartets that involve the hybrid taxon *a* would be considered, and we would erroneously conclude that *abc*|*de* is a cut-edge split on *N*.

## 5. Statistical evidence for cut-edges

### 5.1. Model assumption

The previous section characterizes the presence of cut edges in the full network by that of cut-edges in induced 4-taxon subnetworks. To decide if a 4-taxon semidirected network has a cut-edge split, we judge evidence from quartet count concordance factor data. To achieve this, we strengthen (3.1) in Theorem 3.1.

#### *Assumption* 1.

Consider a gene tree distribution model associated to rooted networks *N* ^+^ on *X*, inducing a probability model *p* on quartet concordance factors. For any *N* ^+^ on *X*, and set *S* = {*a, b, c, d*} ⊆ *X*, with *N*_*S*_ the induced semidirected network on *S*, the following hold: (star) If *N*_*S*_ has a trivial 4-blob (a multifurcation), then

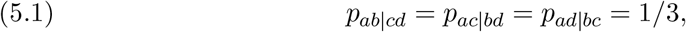

and if *N*_*S*_ does not have a trivial 4-blob, then (5.1) does not hold for generic numerical parameters, and

(T1) If *N*_*S*_ has a cut-edge split *ab*|*cd*, then

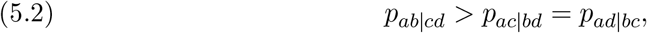

while if *ab*|*cd* is not a cut-edge split of *N*_*S*_, then (5.2) does not hold for generic numerical parameters.

By “generic parameters” in this assumption, we mean those in any set whose complement has measure zero in a measurable parameter space. Here, the edge length parameter associated to a tree edge is naturally in (0, ∞), with Lebesgue measure *µ*. For hybrid edges, where [0, ∞) is the desired parameter interval, treating edges of length 0 as non-generic by the Lebesgue measure is not desirable, as this value is needed to model gene flow between contemporaneous populations. We therefore adopt the measure *δ*_0_ + *µ* on [0, ∞) for hybrid edge lengths. Using *µ* on (0, 1) for inheritance parameters, and the product of these measures on the full numerical parameter space for a network fixes the terminology.

#### Proposition 5.1.

*Let N* ^+^ *be a rooted metric network on X, such that all nodes in non-trivial blobs have degree 2 or 3. Then Assumption 1 holds under the NMSC model, for the restricted numerical parameter space for which N* ^+^ *has no anomalous quartets*.

*Proof*. Hypothesis (T1) is essentially shown in [4, Theorem 1], but that work assumes the network is binary, and uses as its definition of “generic” the Lebesgue measure on [0, ∞) for hybrid edge lengths. We discuss only the necessary adaptations.

For a quarnet induced from *N* ^+^ which has a non-trivial cut-edge split, there can be no multifurcations outside of blobs. That the 2 probabilities in (5.2) are equal follows with no change, while the inequality uses the no-anomalous quartets assumption. For the non-cut edges, a careful reading of the argument shows that at no point is it necessary to rule out edge lengths of 0 for hybrid edges.

For a quarnet with a trivial 4-blob, exchangeability of lineages at that node implies the gene quartet probabilities are uniform. For one with a non-trivial 4-blob, checking the argument of [4, Theorem 1] shows that generically, no two gene quartet probabilities are equal.

We strongly suspect Assumption 1 also holds for the NMSC model with common or correlated inheritance, but do not verify that here. Related matters are investigated in [33], though only in the setting of binary networks, using Lebesgue measure on [0, ∞) to define generic hybrid edge lengths. That work also discusses the Displayed Tree model, for which it is desirable to modify the notion of a quartet CF in the nonbinary setting, as suggested in Section 2, since non-binary nodes in the species network lead to gene tree multifurcations with positive probability.

#### *Remark* 5.2.

Figure 1 provides counter examples showing that Proposition 5.1 does not generally hold for networks with blobs that contain nodes of degree > 3. Both *N*_1_ and *N*_2_ satisfy (5.2) for *ab*|*cd*, yet do not have a cut-edge for that split. Indeed, each of their blobs could be replaced by a single edge of an appropriate positive length *t*_0_, yielding the tree *N*_0_ in Figure 1 with identical quartet CF. (See [8] for a discussion of replacing a biconnected component by a single edge.) Thus some assumption on the network is necessary to ensure that (5.2) provides evidence for a cut-edge. The degree condition within blobs is one such, although it not a necessary one.

### 5.2. Quartet hypothesis tests

Given a collection of gene trees, we apply two hypothesis tests to each quartet count Concordance Factor (qcCF) to test for cut-edge splits of 4-taxon networks. First, the *star test* has null hypothesis that the expected CF frequencies satisfy (5.1). This corresponds to the induced quarnet on *a, b, c, d* having a trivial 4-blob (a multifurcation), possibly with chains of 2-cycles leading to taxa. Failure to reject the null provides evidence of a multifurcation in the network, and hence in the tree of blobs, or inadequate data to obtain a resolution. By default, we use a likelihood ratio test (LRT) for a 3-category multinomial distribution, with 2 degrees of freedom. An alternative option is the standard chi-square test based on the Pearson test statistic and 2 degrees of freedom, asymptotically equivalent to the LRT.

The second test is the T1 test developed in [30] which has

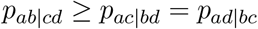

as its null hypothesis. We use this only for the quartet tree *ab*|*cd* that is displayed on the given resolution of the tree of blobs, so rejection of the null gives evidence that the tree of blobs does not display *ab*|*cd*. Note also that we will apply the star test first to test for equality of the expected values. So when the T1 test is applied, one can view it as testing (5.2) with a strict inequality.

## 6. Combining *p*-values

An essential part of our algorithm is combining a number of *p*-values from hypothesis tests applied to CFs from different 4-taxon sets into a single *p*-value. Since these all arise from the same input data, we cannot assume the *p*-values are independent. Moreover, with the network topology unknown, we have no understanding of the exact nature of the non-independence.

Formally, we have *k* null hypotheses *H*_0,*i*_ for 1 ≤ *i* ≤ *k*. For each *H*_0,*i*_, we obtain a *p*-value *p*_*i*_, and then seek a *p*-value to test the global null hypothesis 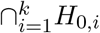. Given a multiset of *p*-values *P* = {*p*_1_, *p*_2_,…, *p*_*k*_}, we consider four methods for combining *p*-values from non-independent tests, providing a correction needed due to multiple testing:

The minimum of the *p*-values, adjusted for the number of tests [40]:

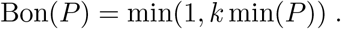

The Cauchy combination *p*-value:

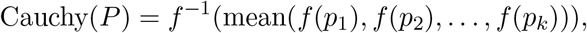

where *f* (*p*) = tan *π*(0.5 − *p*) [26].

The Bon and Cauchy *p*-values combined with Bon [12]:

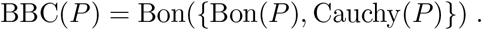

The Bon and Cauchy *p*-values combined with Cauchy [12]:

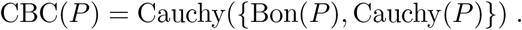

Bon(*P*) is sometimes defined as 1−(1−min(*P*))^*k*^, which gives almost the same value when min(*P*) is small. In the form above, it essentially is the Bonferroni correction. We adopted the name “Bon” rather than “MinP” as used in [12], to avoid the possible misinterpretation that it is simply the minimum of the *p*-values.

The Cauchy combination was more recently introduced, and shown to have good properties under dependency structures where each pair of variables was bivariate normally distributed. While Cauchy does control type-I error, the last two methods, BBC and CBC, were introduced to counteract a loss of power of the Cauchy combination test in certain situations.

The following technical Lemma is needed for a proof of statistical consistency of our algorithms. Informally, this and the probabilistic result of Lemma 8.1 are used to show that a combined *p*-value goes to 0 as the size *n* of a dataset increases, provided at least a single *p*-value tends to 0. The proof is trivial for Bon, but for Cauchy and methods built on it, complications arise since if one *p*-value tends to 1, then the Cauchy combination *p*-value need not go to 0. This is related to the possible lack of power for Cauchy noted by [12]. Thus, we impose an extra condition that *p*-values be bounded away from 1.

### Lemma 6.1.

*Let P* = {*p*_*i*_}_1*≤i≤k*_ ⊂ [0, 1) *denote a multiset of p-values of size k* ≥ 1.

1. *For each method testCo* ∈ {Bon, Cauchy, BBC, CBC} *and k there exists a bounding function B*_*testCo,k*_(*η, D*) *non-decreasing in k, and in its arguments η and D, satisfying where B*_*testCo,k*_(*η, D*) ∼ *ckη for η* → 0 *and D* < 1 *fixed, with c* = 1 *for* Bon, Cauchy *and* CBC, *and c* = 2 *for* BBC.
2. Bon(*P*), Cauchy(*P*), BBC(*P*), CBC(*P*) ≥ min(*P*).

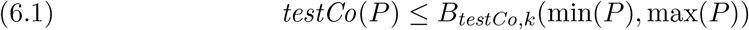

*Proof*. To see part 1, let *m* = min(*P*) and *M* = max(*P*) < 1. First note that Bon(*P*) = *km*, so we can take *B*_Bon,*k*_(*η, D*) = *kη*. For Cauchy, transforming the elements of *P* by the decreasing function *f* (*p*) = tan(*π*(0.5 − *p*)) gives *f* (*m*) and *k* − 1 values between *f* (*M*) and *f* (*m*), whose average is therefore ≥ ((*k* − 1)*f* (*M*) + *f* (*m*)) */k*. Transforming by *f*^*−*1^ then shows (6.1) holds for

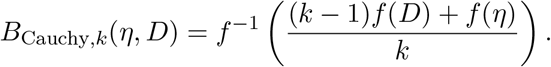

For *D* fixed, *B*_Cauchy,*k*_(*η, D*) ∼ *kη* follows from *f* (*x*) ∼ 1*/*(*πx*) as *x* → 0^+^.

Next, we may take *B*_BBC,*k*_(*η, D*) = 2 min{*B*_Cauchy,*k*_(*η, D*), *B*_Bon,*k*_(*η, D*)} from the definition of BBC, and its claimed properties follow from those for Cauchy and Bon. Finally, since CBC(*P*) ≤ max{Cauchy(*P*), Bon(*P*)} we may choose *B*_CBC,*k*_(*η, D*) = max{*B*_Cauchy,*k*_(*η, D*), *B*_Bon,*k*_(*η, D*)}, which has the claimed properties.

Part 2 is shown easily, using Cauchy(*P*), Bon(*P*) ≥ min(*P*).

## 7. The ECToBlob Algorithm

We present an algorithm, `ECToBlob` (<u>E</u>dge <u>C</u>ontraction for <u>T</u>ree <u>o</u>f <u>B</u>lobs), that begins with any resolution *T*_0_ of the tree of blobs of a network, and successively contracts some of its edges until the data fail to reject a model of gene tree generation on a network with the remaining cut edges. It operates in 3 variants, called `ECToBlob-bi`, `ECToBlob-mul`, and `ECToBlob-quad`. These differ in their use of *p*-values from quartet hypothesis tests for those quartets spanning either <u>bi</u>partitions, current <u>mul</u>tipartitions, or initial <u>quad</u>ripartitions associated to the edges of the resolution of the tree of blobs (Figure 3).

**Figure 3.**
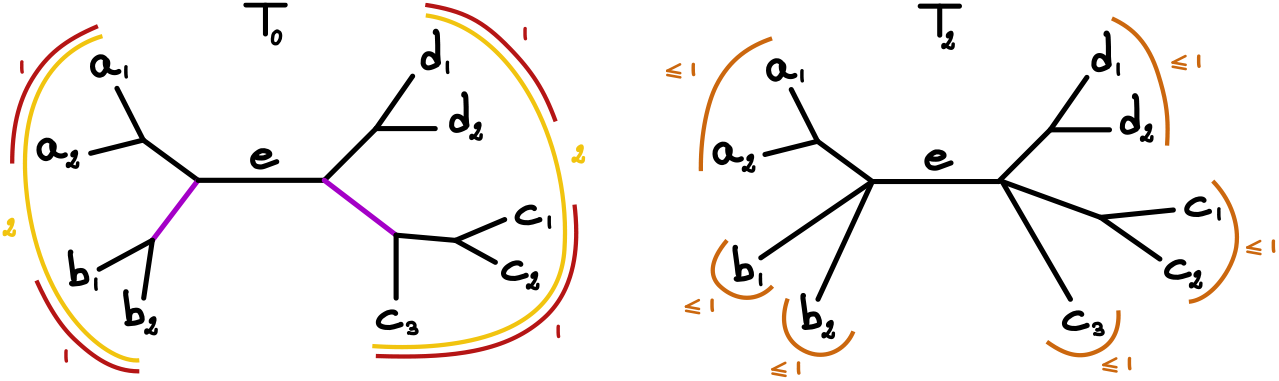
*Example edge e in the starting tree T*_0_ *(left) and in T*_2_ *after contracting 2 edges of T*_0_ *(in purple). To decide whether e should be contracted, we use a collection of quartets. One choice is 𝒮*_bi_(*e*), *from all 4-taxon sets respecting e’s bipartition (yellow). There are* 6 · 10 = 60 *such 4-taxon sets, from choosing 2 taxa in* {*a*_1_, *a*_2_, *b*_1_, *b*_2_} *and 2 in* {*c*_1_, *c*_2_, *c*_3_, *d*_1_, *d*_2_}. *The second is the collection 𝒮* _quad_(*e*) *of 4-taxon sets spanning the quadripartition that e determines in T*_0_ *(red): with 1 taxon in each of* {*a*_1_, *a*_2_}, {*b*_1_, *b*_2_}, {*c*_1_, *c*_2_, *c*_3_} *and* {*d*_1_, *d*_2_}, *thereby using only* (2 *×* 2) *×* (3 *×* 2) = 24 *sets of 4 taxa. A third, 𝒮*_mul_(*e*), *collects the 4-taxon sets spanning the multipartition that e defines in the current tree. In T*_0_, *this is a quadripartition, but to test e in T*_2_, *we instead use the 3 blocks on each side of e, a*_1_*a*_2_|*b*_1_|*b*_2_ *and d*_1_, *d*_2_|*c*_1_|*c*_2_, *c*_3_, *taking at most 1 taxon from each block and at least 2 taxa from each side, here* 5 *×* 8 = 40 *sets of 4 taxa*.

Previously inferred gene trees summarized by quartet count Concordance Factors are input, together with a starting tree, *T*_0_, assumed to be a resolution of the tree of blobs. We suggest using `ASTRAL` to obtain *T*_0_, as justified by Theorem 3.6 and `ASTRAL`’s consistency in inferring a tree optimizing the MQS criterion [29]. However, other heuristics for the same optimization problem may be used, such as `TREE-QMC` [19]. Weighted versions of these methods may also be applicable [42, 9, 31]. While *T*_0_ need not be binary as long as it is a resolution of *T*_*B*_(*N*), for our informal discussion we assume it is. We choose and fix one of the methods from Section 6 for combining possibly dependent *p*-values that will be used throughout the algorithm.

Starting with *T*_0_, we obtain a sequence of trees by successively contracting edges as follows. First we apply, for each qcCF, a test with a star tree as null hypothesis. For each edge *e* in the starting tree *T*_0_, we obtain 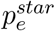, a single star *p*-value for *e* by combining the *p*-values from all sets of 4 taxa from the edge’s quadripartition. Using some choice of test level *β, e* is contracted if we fail to reject the star. Doing so for each edge in *T*_0_, gives a tree 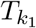, where *k*_1_ is the number of contracted edges. This step contracts edges for which the quartet data does not indicate any *β*-level support for a topology (tree-like or network-like) other than the star. These may be due to true multifurcations, or to inadequate signal in the data (i.e., a soft polytomy, as often occurs in gene tree inference).

Next, for each edge *e* in the current tree *T*_*k*_, we determine a combined edge-specific *p*-value, 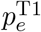 to test the null hypothesis that *e* is in the network’s tree of blobs. This is obtained by combining the T1 test *p*-values across the 4-taxon sets in𝒮 _*qType*_ (*e*) defined in Section 4, where *qType* is `bi, mul` or `quad`. For the `bi` and `quad` quartet types, we use these 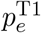 to successively contract an edge, or edges in case of ties, from most weakly supported to strongest supported.For the `mul` type new 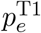 are computed after each contraction, as the current tree becomes more polytomous. Since

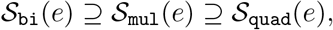

the 3 quartet types use nested subsets of *p*-values with `bi` combining the most and `quad` the fewest. All three approaches yield a sequence of trees 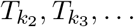 with 0 ≤ *k*_1_ < *k*_2_ < *k*_3_ <…, where *k*_*i*_ indicates the total number of edges of *T*_0_ contracted.

At each step, the tree *T*_*k*_ is assessed in full with a combined T1 *p*-value, *p*_*k*_, found by combining all *p*-values from the T1 test for all resolved quartets that *T*_*k*_ displays. That is, the *p*-values for 4-taxon sets 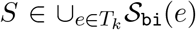 are combined, giving an overall assessment of the remaining cut edges. While the algorithm returns the full sequence 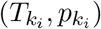, we propose two methods, detailed below, to select the estimated tree of blobs 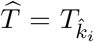 for some *i* in the sequence, based on a test level *α*.

Note that the combined *p*-value assessing all edges remaining in 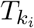 need not monotonically increase with *i*. For instance, if the combination method Bon is used, the combination test for 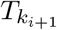 uses fewer *p*-values than that for 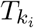, from ignoring the four-taxon sets around the edges in 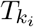that were contracted in 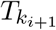. However, at least some of the *p*-values left out of the combination are small, and removing small individual *p*-values and combining fewer *p*-values have opposite impacts on the size of the combined *p*-value. In simulations with true gene trees, the sequence of combined *p*-values assessing 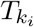was usually increasing, and the rare observed decreases were generally of quite small magnitude. Moreover, we have on occasion observed a similar increase when analyzing empirical data.

We thus propose two selection rules, called *early* and *late*, to select the estimated tree of blobs 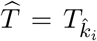. Both select a tree that is not rejected at level *α*, with 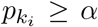. The *early* method simply chooses the first (smallest) 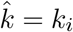 such that 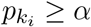. The *late* method chooses the tree from the iteration after the last *p*-value below *α*. If the sequence of combined *p*-values is monotone, then both methods agree. Otherwise, the *early* method tree is more resoluved than the *late* one.

The full procedure is presented more formally in Algorithm 7.1. Note that when *T*_*k*_ is reduced to a star tree, without any internal edges, Step 4 does not have any null hypothesis to test (there are no *p*-values to consider), leading to *p*_*k*_ = 1 and reflecting no evidence to reject the star tree. Therefore, there exists at least one iteration (the last one) with 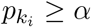, and Step 10 yields well-defined notions of 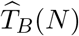.

The time complexity of the ECToBlob algorithm using *qType* = `bi` or `quad` is computed as follows. If |*X*| = *s*, applying the star and T1 test to the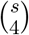 sets of 4 taxa requires time 𝒪 (*s*^4^). Then, for each of the at most *s* − 3 internal edges of a tree *T*_0_, determining which *p*-values must be combined for these tests and combining them is also 𝒪 (*s*^4^), for a total of 𝒪 (*s*^5^). Sorting the individual *p*-values is negligible in comparison. At each of the at most *s* − 3 edge contracting steps, we must compute a combined *p*-value for the tree, requiring time at most 𝒪 (*s*^4^), or 𝒪 (*s*^5^) in total. This gives time complexity 𝒪 (*s*^5^) for the entire algorithm.

For *qType* = `mul` there is additional computation as new combined *p*-values must be found at each contraction step, for all edges incident to the one contracted. For a binary *T*_0_ the *k*th edge contracted might be incident to as many as 4 + *k* other internal edges, so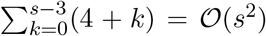 new edge computations may be needed with total time𝒪 (*s*^6^). This gives a total time complexity 𝒪 (*s*^6^).

Our implementation of a function `ECToBlob` in the R package `MSCquartets` has default values of *β* = 0.80 and *α* = 0.05. The large value of *β* means that few edges are contracted at the first iteration, only those with CFs very close to 1/3.

In addition to 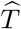, the function returns additional values, so that a different choice of test level *α* and/or selection rule require no additional computation. Specifically, it returns the full sequence of trees *T*_0_, 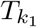, 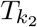,…, and for each *T*_*k*_, both its overall *p*-value *p*_*k*_ and its edgespecific *p*_*e*_-values from the corrected tests (star for *T*_0_, T1 for later trees). Finally, the output includes the index *i* of the inferred tree of blobs 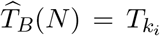 selected at level *α* by both *early* and *late* criteria. An option for plotting the full sequence of trees allows for the easy sequential inspection of each tree *T*_*k*_ and their changing overall *p*-values.

### Algorithm 7.1

`ECToBlob`

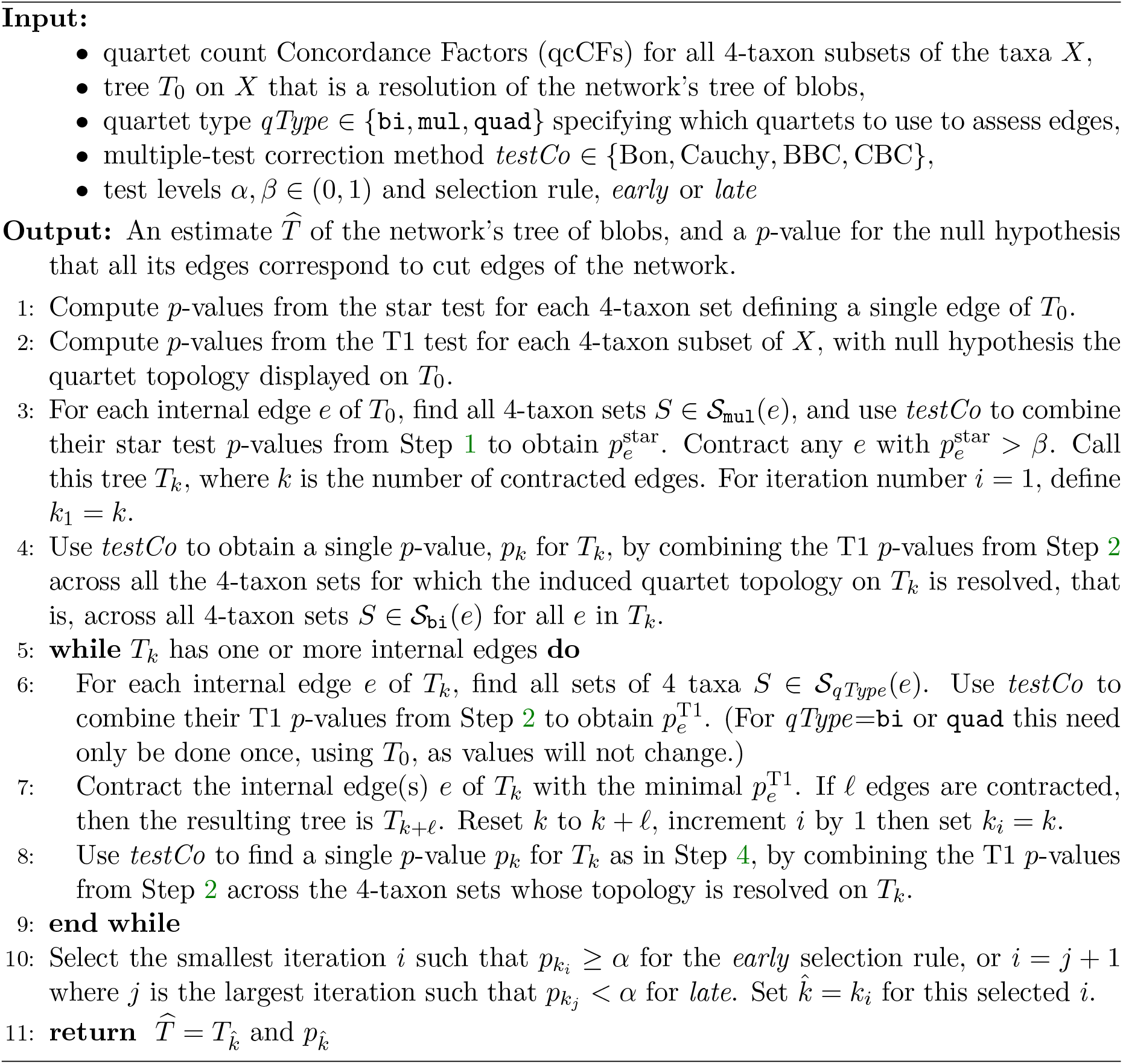

## 8. Statistical consistency

Since the `ECToBlob` algorithm is built on hypothesis tests, to show consistency we need to establish that it draws the correct conclusions from these, with a probability that approaches 1 with more and more gene trees. To ensure this, it is necessary to adjust the levels of the tests as the dataset size increases, simultaneously controlling the probability of both correct rejection and correct failure to reject of their null hypotheses.

To this end, we use the following lemma, as well as the bounds on the combinations of *p*-values established in Lemma 6.1. Note that we do not use any results from [26], which introduced the Cauchy combination test, since its bivariate normal assumption on pairs of *p*-values may not apply here.

The star test is a likelihood ratio test (by default, in our implementation) with a standard 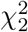 test distribution, and the T1 test uses a likelihood ratio statistic with a test distribution which, except at the boundary point (1/3,1/3,1/3), is asymptotically a 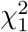 distribution [30]. From well-known properties of likelihood ratio tests, if the expected quartet CFs satisfy the null hypothesis, then the *p*-value from each individual 4-taxon set has a uniform distribution on [0, 1], asymptotically as the number of genes increases. If instead the null hypothesis is false, then the power approaches 1 with increasing number of gene trees.

While the behavior of individual 4-taxon tests is well understood, we need the following result, along with Lemma 6.1, to address the behavior of combined *p*-values.

### Lemma 8.1.

*Let K* ≥ 1 *be fixed. Consider a sequence of sets of k* ≤ *K possibly dependent random variables*

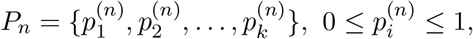

*where as n* → ∞ *each* 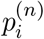 *is individually asymptotically uniformly distributed. Then for any ϵ* > 0 *there exist* 0 < *δ*< 1*/*2 *and M such that for n* > *M*,

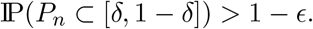

*Proof*. Using the union bound, note that

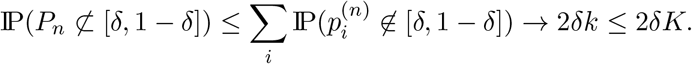

We may assume *ϵ* ≤ 1. Letting *δ*= *ϵ/*4*K*, so 2*δK* = *ϵ/*2, the existence of such an *M* follows.

### Theorem 8.2.

*There exist sequences* {*α*_*n*_} *and* {*β*_*n*_}, *independent of the network N* ^+^, *such that for any resolution T*_0_ *of T*_*B*_(*N*^+^), *any variant of the `ECToBlob` algorithm, and any model meeting Assumption 1, using α*_*n*_, *β*_*n*_, *and qcCFs from a sample of n unrooted topological gene trees, the output* 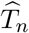*satisfies*

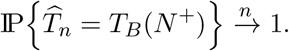

*Proof*. Consider the sequences defined by

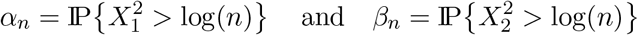

where 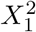 and 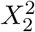 follow a 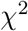 distribution with 1 and 2 degrees of freedom respectively. Then 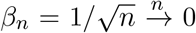, and 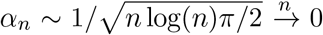. Let *s* be the number of taxa on *N*^*+*^, and let *ϵ* > 0. We will prove that for *n* sufficiently large, with probability 1 − *ϵ*, all the various combination tests computed by the algorithm correctly indicate whether edges of *T*_0_ should be contracted or not.

We begin with the star test. With our assumption of generic numerical parameters, we seek to find *M*_star_ such that for each edge *e* of *T*_0_, the combined star test *p*-value using the 4-taxon sets in *S*_mul_(*e*) will, with probability > 1 − *ϵ/*(2*s*), fail to reject the null hypothesis at level *β*_*n*_ for *n* ≥ *M*_star_ if the expected CF is (1*/*3, 1*/*3, 1*/*3) for all such 4-taxon sets, and reject it otherwise. Since *T*_0_ has at most *s* − 3 internal edges this will ensure the star test will make the correct conclusion for all these edges simultaneously with probability > 1 − *ϵ/*2.

To this end, suppose first that all sets in *S*_mul_(*e*) have expected CFs of (1*/*3, 1*/*3, 1*/*3). As the null star hypothesis is met, each individual *p*-value is uniformly distributed as *n* → ∞.

### Lemma 8.1

then implies that there are 1*/*2 > *δ*> 0 and 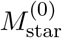such that if 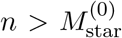thenwith probability > *q*_*ϵ*_ = 1 − *ϵ/*(4*s*) all individual *p*-values satisfy *δ*≤ *p* ≤ 1 − *δ*. Now, let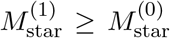 be such that 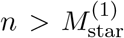 implies *β*_*n*_ < *δ*. Then for 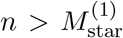, all individual *p*-values satisfy *β*_*n*_ < *p* 1 *δ*with probability > *q*_*ϵ*_. By Lemma 6.1 item 2, the combined *p*-value for *e* is 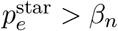 with probability > *q*_*ϵ*_.

Now consider an edge *e* for which one or more of the expected CFs is not (1*/*3, 1*/*3, 1*/*3). For the expected CFs that are (1*/*3, 1*/*3, 1*/*3) we have already shown there is a 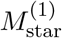 such that when 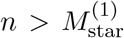 the *p*-values are in (*β*_*n*_, 1 − *δ*] with probability > *q*_*ϵ*_. The remaining 4-taxon sets, for which the alternative hypothesis is true (the CF is not (1*/*3, 1*/*3, 1*/*3)), form 𝒮^*a*^(*e*) ⊂ *S*_mul_(*e*). Let *η*_*n*_ > 0 be such that the bound from Lemma 6.1 item 1, satisfies 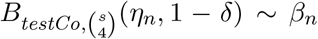, where *testCo* is the test correction method used to combine *p*-values. Let *b*_*n*_ = −2 log(*η*_*n*_), so that IP 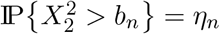 and an individual *p*-value is below *η*_*n*_ exactly when the test statistic is above *b*_*n*_.

### Lemma 6.1

then implies *b*_*n*_ ∼ log *n*. For *S* ∈ 𝒮^*a*^(*e*), let 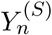be its test statistic for the star test from *n* gene trees. Since the null star hypothesis is false on *S*, it is well known for the likelihood ratio test statistic 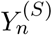 that 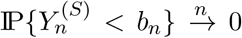 if *b*_*n*_*/n* → 0 [35, §6.4.2 for example]. (If the test uses the Pearson chi-square statistic instead, the same result holds [35,§6.5.4, for example].)

Therefore, 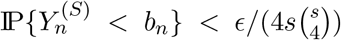 for all *n* > *M*_star,*S*_, for some *M*_star,*S*_. Define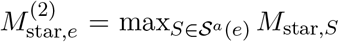. Since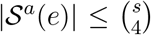, we have with probability at least *q*_*ϵ*_, that 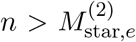implies 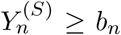 for all *S* ∈ 𝒮^*a*^(*e*). In other words, with probability ≥ *q*_ε_, the *p*-value from each *S* ∈ 𝒮^*a*^(*e*) is below *η*_*n*_. Then when 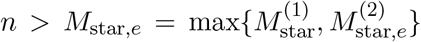,the minimum individual *p*-value for *e* is at most *η*_*n*_ and the maximum individual *p*-value is at most 1 − *δ*, and by Lemma 6.1 item 1, the combined 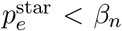 with probability > 1 − *ϵ/*(2*s*).

Thus there exist *M*_star_ = max_*e*_{*M*_star,*e*_} such that, with probability > 1 − *ϵ/*2, for any *n* > *M*_star_, 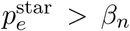 if all 4-taxon sets defining edge *e* have CFs of (1*/*3, 1*/*3, 1*/*3), and 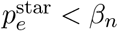 otherwise.

The T1 tests are handled similarly. The quartet types `bi`, `mul`, and `quad` determine different ensembles of 4-taxon sets for each edge, to which the T1 test is applied. At a given stage of edge contraction, consider the set of 4-taxon sets 𝒮^*a*^(*e*) of the chosen quartet type for which the T1 null hypothesis is false. Then by Corollaries 4.2 and 4.3, 𝒮^*a*^(*e*) is empty exactly when *e* is a cut-edge of *N* ^+^. Arguing as for the star test, there exists an *M*_T1_ such that with probability > 1 − *ϵ/*2 for any *n* > *M*_T1_, the combined *p* value 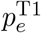 for an edge *e* of *T*_*k*_ is in (*α*_*n*_, 1 − *δ*) if *e* is a cut-edge of *N* ^+^, and 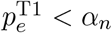 otherwise. This second outcome correctly determines splits not on *T*_*B*_(*N* ^+^).

With the *α*_*n*_, *β*_*n*_ as above and *M* = max{*M*_star_, *M*_T1_}, we have the following, with probability > 1 − *ϵ*: the star tests in the algorithm correctly determine the edges in *T*_0_ that (erroneously) resolve multifurcations in the network, and the T1 tests correctly determine which edges in *T*_0_ are consistent with cut-edge splits in the network. In Step 3 of the `ECToBlob` algorithm 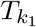 is obtained from *T*_0_ by contracting precisely those edges that resolve multifur-cations in *T*_*B*_(*N* ^+^) (with probability > 1 − *ϵ/*2). In repeated iterations of Step 5, edges not in *T*_*B*_(*N* ^+^) will all have *p*-values below *α*_*n*_ and edges in *T*_*B*_(*N* ^+^) will all have *p*-values above *α*_*n*_, hence edges not in *T*_*B*_(*N* ^+^) will be contracted first (with probability > 1 − *ϵ/*2).

Finally, we consider the overall *p*-value *p*_*k*_ associated with each tree *T*_*k*_ in the sequence. The *early* and *late* selection will both select the correct tree if *p*_*k*_ < *α*_*n*_ whenever *T*_*k*_ retains an edge that is not a cut-edge in *N* ^+^, and *p*_*k*_ > *α*_*n*_ otherwise. This occurs with probability at least 1 − *ϵ/*2 using our prior choice of *M*, because *p*_*k*_ combines the T1 test *p*-values from 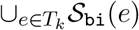, of size at most 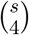. If all edges *e* in *T*_*k*_ are cut-edges in *N* ^+^, then all the individual *p*-values are in (*α*_*n*_, 1 − *δ*) and *p*_*k*_ > *α*_*n*_. Otherwise, at least one input *p*-value is < *η*_*n*_ and *p*_*k*_ < *α*_*n*_ by the same argument as before. Thus 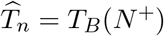 with probability > 1 − *ϵ*.

While specific choices for *α*_*n*_ and *β*_*n*_ were made in our proof, others could be used instead. The choice for *α*_*n*_ above corresponds to the BIC [34], as the T1 test uses 1 degree of freedom. For the star test, with 2 degrees of freedom, the BIC corresponds to comparing the likelihood ratio statistic to 2log(*n*) rather than the choice log(*n*) here, that is, a test level *β*_*n*_ = 1*/n* instead of 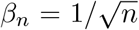 used in the proof. Any valid choice needs *α*_*n*_, 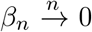 in order to fail to reject a true null hypothesis consistently. However, we also need *o*(*n*) cutoff values (e.g., log(*n*) in the proof), to ensure they grow slower than the non-centrality parameter of each non-null hypothesis, so that a false null hypothesis is rejected with increasing probability. Future work could focus on an optimal choice, which might depend on specific details such as the number of taxa or number of individual tests to combine for each edge. Different costs for the 2 types of errors, false discoveries (failing to contract an edge not in *T*_*B*_(*N* ^+^)) and missed edges (contracting an edge that truly is in *T*_*B*_(*N* ^+^)), could also be incorporated.

Note that with empirical datasets, the star test may lead to the contraction of true network edges if the data provides little support for any resolution. That is, a multifurcation in an inferred tree of blobs may represent not a blob, but rather a ‘soft’ polytomy, expressing inadequacy of the data to obtain a valid resolution. A similar effect may produce multifurcations in which one or more true contracted blobs may merge with adjacent soft polytomies to give a multifurcation of greater degree than the true one.

### *Remark* 8.3.

The argument in the proof of Theorem 8.2 is easily adapted to strengthen the consistency results for the NANUQ [1] and TINNiK [5] algorithms, which use quartet hypothesis tests in a similar way (though without combining *p*-values). This makes explicit some valid choices of sequences of test levels for those algorithms.

### *Remark* 8.4.

If `ECToBlob` is given an input tree *T*_0_ that is not a resolution of the tree of blobs, the argument above indicates it would still contract any edges of *T*_0_ that do not define cut-edge splits in the true network. That is, asymptotically it should return a “coarsened” version of the tree of blobs, so that the cut edges in its output still represent cut edges of the network, although the network’s cut edges absent from *T*_0_ remain absent.

## 9. Data analysis

We performed simulations to assess the accuracy and speed of `ECToBlob` under idealized circumstances, and applied `ECToBlob` to a genomic dataset of Neotropical cats [25]. Summaries of these results are given below, with additional figures and full simulation settings and outcomes given in Supplementary Materials A.

### 9.1 ECToBlob accuracy in simulations

Using `PhyloCoalSimulations` [16, v1.1.0], we simulated *m* = 300, 500, 1000 gene trees on a model network *N* on 30 taxa (shown in Figure S1 of Supplementary Materials A). This network has 5 reticulations forming a 4-blob and two 7-blobs. Branch lengths for *N* were drawn from the Γ(20, 1*/*20) distribution, with inheritance parameters *γ* varying to make inference more challenging. *N* was then scaled by factors *s* = 1, 0.75, 0.5, 0.25 to give increasing ILS levels. For each value of *m* and *s*, 100 replicate datasets were generated then analyzed with `ECToBlob` using the ASTRAL tree for *T*_0_ and all 12 combinations of quartet type (`bi, mul, quad`) and multiple test correction method (Bon, BBC, Cauchy, CBC). Default test levels *β* = 0.8 and *α* = 0.05 were used, under both the *early* and *late* selection rule. The Robinson Foulds (RF) distances from the `ECToBlob` tree to the true tree of blobs *T*_*B*_(*N*) were computed to measure the topological accuracy.

The RF distances, averaged across replicates, are shown in Figure 4 for the *late* selection rule. Under these settings, the test corrections Bon and BBC substantially outperformed the Cauchy and CBC variants. Using Bon or BBC with any *qType*, `ECToBlob` accuracy was excellent, with near-perfect recovery of the true tree of blobs under moderate to moderately high ILS (*s* ≥ 0.75). Accuracy degraded with high ILS (*s* = 0.25) or few gene trees (Figure 4 and Table S1 of Supplementary Materials A). Similar results from other simulations suggest Bon and BBC should be preferred options for analyses.

**Figure 4.**
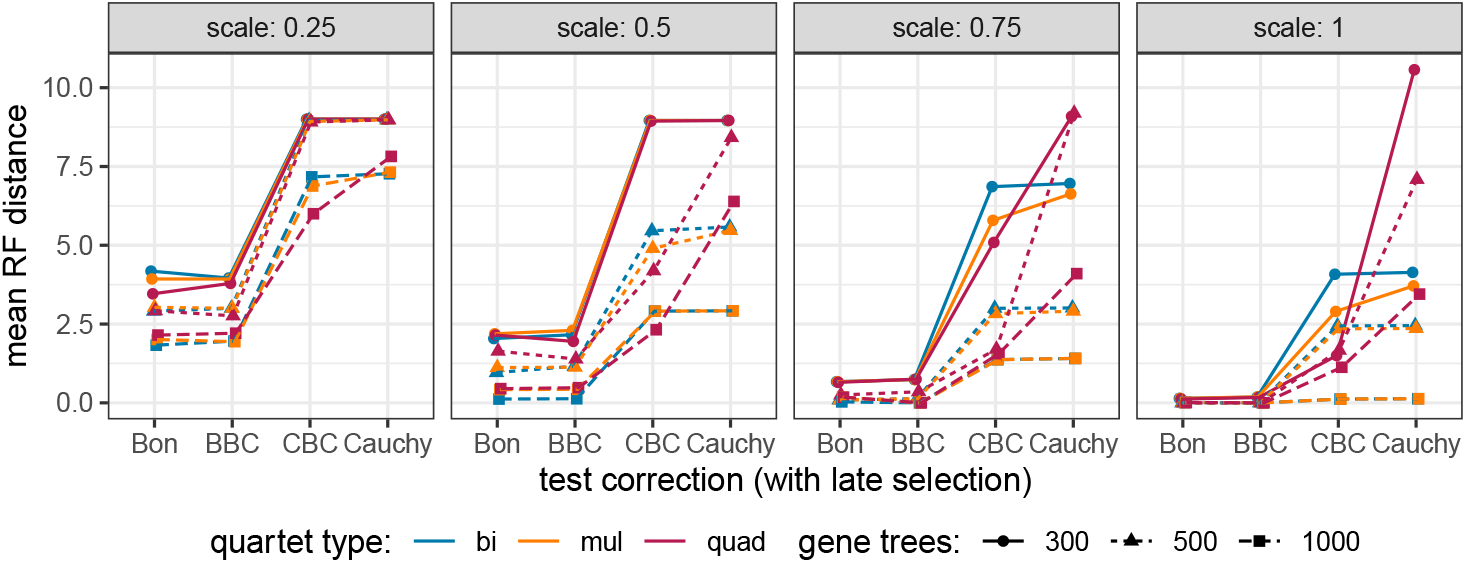
Mean RF distance to true tree of blobs T_B_ (N). The Bon and BBC corrections perform better on these simulated data. Results using the early selection rule are essentially identical.

Computational time was primarily spent tallying quartet CFs from gene trees and computing *p*-values for individual quartet hypothesis tests. These computations, however, need only be performed once, and can then stored for additional ECToBlob or other downstream analyses. Thus Table 1 reports timing information in two categories: gene tree processing (read gene trees, compute quartet hypothesis test *p*-values) and `ECToBlob` tree construction. Mean gene tree processing time is reported for each value of *m* and averaged over 400 datasets (100 replicates for each *s*), while `ECToBlob` timing, which is independent of *m*, is averaged over 1200 replicates (100 replicates for each *m, s*), since it varies little with *s*.

**Table 1.**
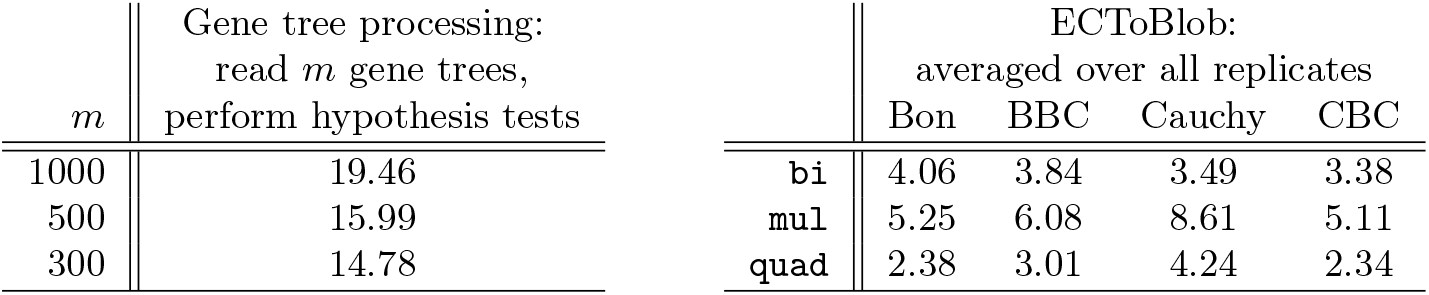
*Time (in secs) to compute the* `ECToBlob` *tree, averaged over datasets. Analyses were performed with a Macbook Pro M3 chip with 36 Gb memory*.

### 9.2 ECToBlob analysis of Leopardus

Controversy remains in regard to the phylogeny of Neotropical cats. As part of their whole-genome analyses study, Lescroart et. al. [25] investigated taxonomic relationships, including introgression and hybrid speciation events, between 16 taxa. These filtered data, 16338 local trees obtained using maximum likelihood from alignments of 100 kb non-overlapping genomic fragments, were analyzed with multiple methods, display significant gene tree discordance, and support a hypothesis of reticulate evolutionary history. We applied `ECToBlob` to these local trees, supplying the ASTRAL tree (Box 1, right) for the starting tree *T*_0_, setting test levels *β* = 0.95 and *α* = 0.05. Selection rules *early* and *late* returned the same trees.

All 12 combination methods returned the tree sequence *T*_0_, *T*_4_, *T*_5_, *T*_6_ (shown in Figure 5 below), followed by other trees. At level *α* = 0.05 the Bon and BBC methods selected *T*_6_ whereas the Cauchy and CBC methods chose *T*_5_.

**Figure 5.**
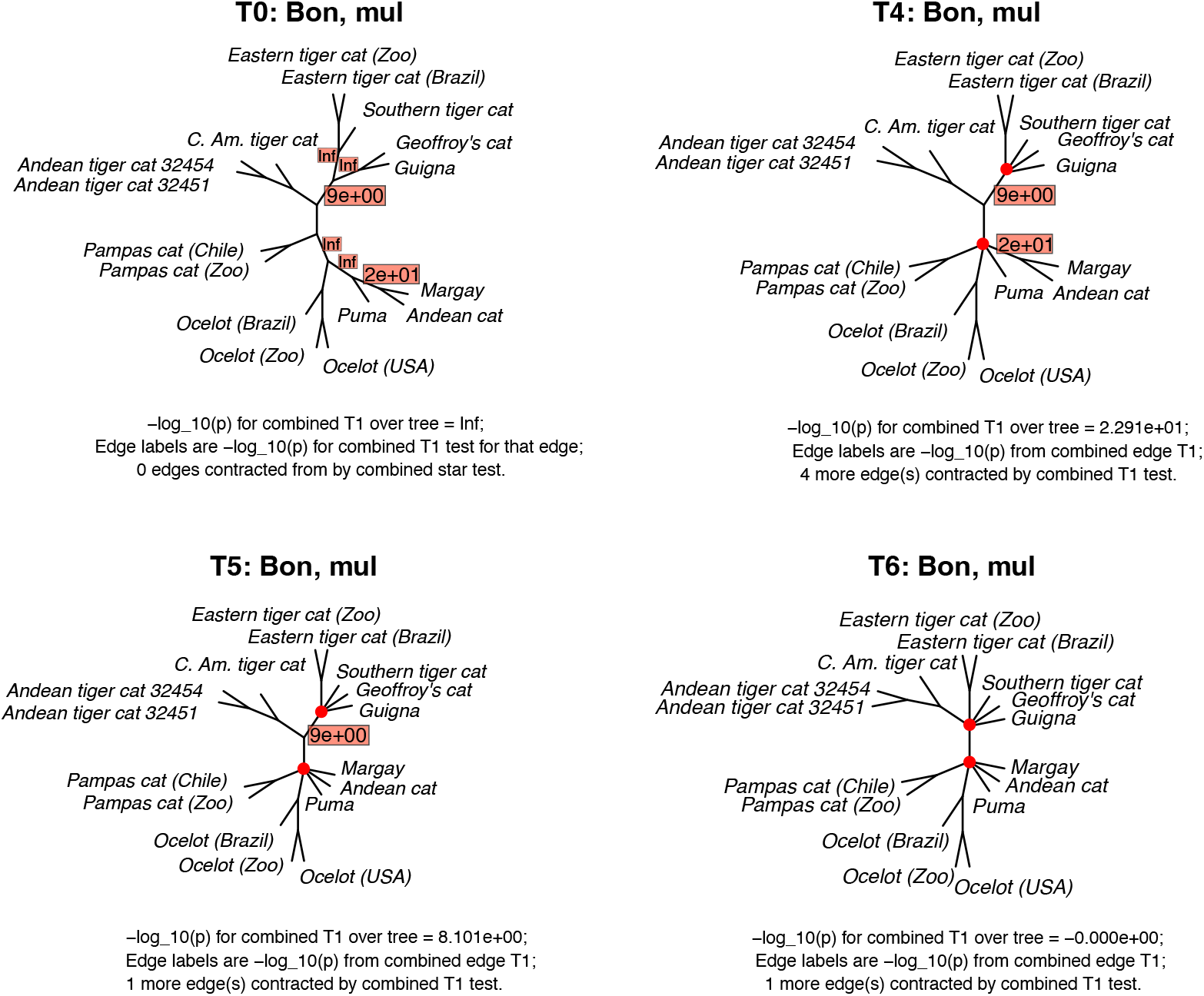
*ECToBlob sequence of trees T*_*k*_ *for all qTypes, starting with the ASTRAL tree (top left). The trees are annotated with their p-values (shown in boxes as*− log_10_(*p*)*) from the mul quartet type and Bon multiple-test correction. At level α* > 10^−7^, *including α* = 0.05, *ECToBlob returns the tree T*_6_ *(bottom right)*.

Interestingly, these both differ from that inferred by TINNiK in [5], which is *T*_4_. As seen in *T*_4_ and *T*_5_, the combined edge *p*-values for the edges contracted to give *T*_6_ are rather small, on the order of 10^−20^ and 10^−9^, showing poor support as cut edges. Moreover, the sequence of combined *p*-values for the trees *T*_0_, *T*_4_, *T*_5_, *T*_6_ is approximately 0, 10^−23^, 10^−8^, 1, giving strong evidence to select *T*_6_ at any test level larger than 10^−7^.

Note that for empirical data a researcher might deviate from the default values for *α, β*, since inferred local trees are likely to differ from a valid sample under the NMSC model. Decreasing *α*, which generally reduces the number of edges contracted, may be a desirable way to address such concerns. However, domain expertise is needed to make such judgments, so we do not do so here.

## 10. Discussion

### Comparison with TOB-QMC

Although following similar outlines, our work here on the `ECToBlob` algorithm differs in several ways from that presented for `TOB-QMC` in [13].

First, our method uses the T1 test, which tests a particular quartet tree topology, rather than the T3 test used by TOB-QMC, which has as its null hypothesis any of the 3 resolved topologies. The use of the T1 test for blobs of arbitrary level is based on the combinatorial results of Theorem 4.2 and Theorem 4.3, and we know of no proof of a stronger result that would justify using the T3 test. Thus the formal justification for `TOB-QMC` is currently limited to level-1 networks [13, Theorem 2], although simulations suggest it may perform well beyond that case. However, since the null hypothesis for T1 is nested in that for T3, the T1 test has greater power to give evidence that an edge should be contracted, so its use may be preferred.

Second, rather than using a single quartet *p*-value as a basis for collapsing an edge, we use a collection of them, and a statistically justifiable procedure to account for the multiple testing of non-independent hypotheses that entails. This is particularly important since dif-ferent edges have different numbers of quartets associated to them, and are thus potentially susceptible to differing type-1 error rates if a small single-test *p*-value occurs simply by chance. See Supplementary Materials B for simulations indicating that improved performance can be achieved by `ECToBlob` in some circumstances, which we attribute to its multiple testing corrections. While `ECToBlob` may be limited to more moderate sized datasets than `TOB-QMC`, its processing time for 50 taxa is a few minutes, and for 100 taxa may be only several hours (See Supplementary Materials A). We believe that `ECToBlob`’s loss of speed compared to `TOB-QMC` is worthwhile, as it results from its strengthened statistical methodology.

We also consider the contraction of edges in steps, rather than all at once based on a single pre-chosen test level, to be important. The resulting putative tree of blobs should not be judged by the *p*-values associated to edges that are contracted, but rather by those remaining. Our goal is to find the most resolved tree that cannot be rejected by a global test of all remaining cut edges. While this iterative contracting process is also slower than `TOB-QMC`’s approach, again, we maintain that it provides greater statistical validity.

Finally, we think that scientists will find the ability to easily view the succession of edge contractions useful in practice. In many cases, multifurcations form at different steps in the sequence, suggesting which blobs are most strongly supported, and those for which evidence is more marginal. With a sequence of trees returned, more insight can be drawn.

#### Comparison with TINNiK

`TINNiK` [5], the first method developed for tree of blobs inference, uses hypothesis testing in its initial step to determine induced quarnets of the unknown network that have 4-blobs. Then, a combinatorial inference rule is applied to expand the set of 4-taxon sets defining each blob to include some that were initially (and correctly) judged not to have 4-blob quarnets. It makes no correction for multiple testing, and the combinatorial rule implicitly assumes all test conclusions are correct. While `TINNiK` was shown to be statistically consistent, we expect that the correction for multiple testing of `ECToBlob` will give it better performance across a wide range of datasets. However, since `TINNiK`’s procedure for finding a tree of blobs is quite different, and in particular, is not based on first finding a resolution of the tree of blobs to serve as an initial tree, it provides a useful alternative perspective as methods for tree of blobs inference continue to progress.

#### Further developments and caveats

As proposed, the overall *p*-value *p*_*k*_ to assess the fit of *T*_*k*_ is obtained by combining the T1 *p*-values from the 4-taxon sets whose topology is resolved in *T*_*k*_. Alternatively, one may consider combining the edge *p*-values, *p*_*e*_, across all edges *e* in *T*_*k*_. This approach would have the advantage of combining fewer input *p*-values, which may increase power. A downside is that some individual quartet *p*-values would be indirectly used multiple times, as some 4-taxon sets may be used to derive *p*_*e*_ for multiple edges *e*. Future work could study this modification, perhaps with a non-uniform weighting of the edge-specific *p*_*e*_ values for the Cauchy combination test.

While we have focused here on using qcCFs to test the topology of each 4-taxon set, our framework could easily be extended to other data types and statistical tests, since at its core `ECToBlob` takes quartet *p*-values as input. The input star test *p*-values could come from any valid test of a multifurcation, and the input T1 test *p*-values from any test that the 4-taxon subnetwork has the desired edge split.

For example, input data could come from a concatenated multiple sequence alignment, summarized by pairwise distances *D* between taxa, instead of quartet CFs. Indeed, tree-like distances can be characterized by the 4-point condition, which involves 3 sum-of-pair values similar to the 3 concordance factors: (SP_*ab*|*cd*_, SP_*ac*|*bd*_, SP_*ad*|*bc*_) where SP_*xy*|*zt*_ = *D*(*x, y*) + *D*(*z, t*). Under adequate conditions on the network and on the model, the analogue of Assumption 1 for distances is met [33], and the tree of blobs is identifiable from average pairwise distances. The `ECToBlob` framework could be applied to this input data type, under models for which a trivial 4-blob (star) corresponds to equal SP values, and a cut-edge split (T1) corresponds to SP_*ab*|*cd*_ < SP_*ac*|*bd*_ = SP_*ad*|*bc*_, and provided that an adequate test is designed for each. A similar approach might use the Log-det distance, drawing on results of [6] for ultrametric networks.

As described here, the same genomic data are used to estimate *T*_0_ and to estimate the quartet counts used by `ECToBlob`. However, using the same data twice can lead to biases [39]. For example, consider a 4-taxon set *S* whose subnetwork is tree-like, but whose split is not a cut-edge split of *N* ^+^. Then the T1 test *p*-value for *S* has a uniform distribution in (0, 1) (asymptotically). However, conditional on its split estimated to be in *T*_0_, this distribution may be non-uniform. Data splitting is a standard way to avoid such a bias, e.g., using *m*_1_ genes to infer *T*_0_, and the other *m*_2_ = *m*−*m*_1_ genes to obtain qcCFs and *p*-values to test edges in *T*_0_. In practice, though, these smaller datasets can result in more estimation error in *T*_0_, and less power for each test. Future work could study this issue and strategies to avoid bias, perhaps with the derivation of post-selection *p*-values conditional on the selection of edges in *T*_0_, as in selective inference (see e.g. [38] for LASSO-type methods, [23] for a review).

Finally, it would be desirable to drop the “no anomalous quartet” assumption we have made for `ECToBlob`. This could easily be done for our use of the T1 test, along the lines of how the cut test of [5], can serve as a analog of the T3 test when that assumption is not made. Nonetheless we still need the assumption to justify that the MQS criterion gives a resolution of the tree of blobs. Indeed, without that assumption, the correct quartet topology need not be indicated by the largest entry of the expected CF, but rather by the CF entry that differs from the others. Developing an algorithm to give a resolution of the tree of blobs based on that fact would be valuable. Of course, the `TINNiK` algorithm, which does not require such a resolution, provides consistent inference even in the face of anomalous quartets, although a thorough comparison of performance is still needed.

## Acknowledgments

This research began while the authors were in residence at the Institute for Computational and Experimental Research in Mathematics in Providence, RI, during the “Theory, Methods, and Applications of Quantitative Phylogenomics” program in Fall 2024, supported by National Science Foundation grant DMS-1929284. We thank O. Gasçuel, K. Wicke, and N. Holtgrefe for their contributions in initial discussions, and J. Dai, Y. Han, and E. Molloy for sharing their pre-publication work on `TOB-QMC`.

## Author contributions

All authors made substantial intellectual contributions to the work, including conceptualization, theory, software development, and simulation study. All authors participated in writing the manuscript and approved its final form.

## Data & code availability

The R package implementing `ECToBlob, MSCquartets` v. 3.3, is available at https://cran.r-project.org/package=MSCquartets. Code to reproduce simulated data and ECToBlob analyses is at https://github.com/eallman/2026_ECToBlob_Sims. A vignette showing the usage of the implementation can be found at https://cran.r-project.org/web/packages/MSCquartets/vignettes/ECToBlob.html.

## REFERENCES

[1] E. Allman, H. Baños, and J. Rhodes, NANUQ: A method for inferring species networks from gene trees under the coalescent model, Algorithms Mol. Biol., 14 (2019), pp. 1–25, 10.1186/s13015-019-0159-2.

[2] E. Allman, H. Baños, J. Mitchell, and J. Rhodes, The tree of blobs of a species network: Identifiability under the coalescent, J. Math. Biol., 86 (2023), p. 10, 10.1007/s00285-022-01838-9.

[3] E. S. Allman, C. Ané, H. Baños, and J. A. Rhodes, Beyond level-1: Identifiability of a class of galled tree-child networks, Bulletin of Mathematical Biology, 87 (2025), p. 166, 10.1007/s11538-025-01545-8.

[4] E. S. Allman, H. Baños, J. D. Mitchell, and J. A. Rhodes, The tree of blobs of a species network: identifiability under the coalescent, Journal of Mathematical Biology, 86 (2022), p. 10, 10.1007/s00285-022-01838-9.

[5] E. S. Allman, H. Baños, J. D. Mitchell, and J. A. Rhodes, TINNiK: inference of the tree of blobs of a species network under the coalescent model, Algorithms for Molecular Biology, 19 (2024), p. 23, 10.1186/s13015-024-00266-2.

[6] E. S. Allman, H. Baños, and J. A. Rhodes, Identifiability of species network topologies from genomic sequences using the logdet distance, Journal of Mathematical Biology, 84 (2022), p. 35, 10.1007/s00285-022-01734-2.

[7] E. S. Allman, H. Baños, J. A. Rhodes, and K. Wicke, NANUQ+: A divide-and-conquer approach to network estimation, Algorithms for Molecular Biology, 20 (2025), p. 14, 10.1186/s13015-025-00274-w.

[8] C. Ané, J. Fogg, E. Allman, H. Baños, and J. Rhodes, Anomalous networks under the multispecies coalescent: theory and prevalence, J. Math. Biol., 88 (2024), 10.1007/s00285-024-02050-7.

[9] E. Avni, R. Cohen, and S. Snir, Weighted quartets phylogenetics, Systematic Biology, 64 (2014), pp. 233–242, 10.1093/sysbio/syu087.

[10] H. Baños, Identifying species network features from gene tree quartets, Bul. Math. Biol., 81 (2019), pp. 494–534, 10.1007/s11538-018-0485-4.

[11] M. B. Bjorner, E. K. Molloy, C. N. Dewey, and C. Solis-Lemus, Detectability of varied hybridization scenarios using genome-scale hybrid detection methods, Bulletin of the Society of Systematic Biologists, 3 (2024), 10.18061/bssb.v3i1.9284.

[12] Z. Chen, Robust tests for combining p-values under arbitrary dependency structures, Scientific Reports, 12 (2022), p. 3158, 10.1038/s41598-022-07094-7.

[13] J. Dai, Y. Han, and E. K. Molloy, Quartet-based species tree methods enable fast and consistent tree of blobs reconstruction under the network multispecies coalescent, bioRxiv, version 5 (2026), 10.1101/2025.11.05.686850.

[14] V. Dinh and H. Baños, Misspecification strikes: Astral can mislead in the presence of hybridization, even for nonanomalous scenarios, Molecular Biology and Evolution, 42 (2025), p. msaf049, 10.1093/molbev/msaf049.

[15] A. K. Englander, M. Frohn, E. Gross, N. Holtgrefe, L. van Iersel, M. Jones, and S. Sullivant, Identifiabiltiy of phylogenetic level-2 networks under the Jukes-Cantor model, bioRxiv, version 2 (2025), 10.1101/2025.04.18.649493.

[16] J. Fogg, E. S. Allman, and C. Ané, PhyloCoalSimulations: A simulator for network multispecies coalescent models, including a new extension for the inheritance of gene flow, Systematic Biology, 72 (2023), pp. 1171–1179, 10.1093/sysbio/syad030.

[17] M. Frohn, N. Holtgrefe, L. van Iersel, M. Jones, and S. Kelk, Reconstructing semi-directed level-1 networks using few quarnets, Journal of Computer and System Sciences, 152 (2025), p. 103655, 10.1016/j.jcss.2025.103655.

[18] M. Habib, K. Roy, S. Hasan, A. H. Rahman, and M. S. Bayzid, Terraces in species tree inference from gene trees, BMC Ecology and Evolution, 24 (2024), 10.1186/s12862-024-02309-z.

[19] Y. Han and E. K. Molloy, Improving quartet graph construction for scalable and accurate species tree estimation from gene trees, Genome Research, 33 (2023), pp. 1042–1052, 10.1101/gr.277629.122.

[20] M. R. Haque and L. Kubatko, A global test of hybrid ancestry from genome-scale data, Statistical Applications in Genetics and Molecular Biology, 23 (2024), p. 20220061, 10.1515/sagmb-2022-0061.

[21] M. Hill, B. Legried, and S. Roch, Species tree estimation under joint modeling of coalescence and duplication: sample complexity of quartet methods, Annals of Applied Probability, 32 (2022), pp. 4681–4705, 10.1214/22-AAP1799.

[22] K. T. Huber, L. v. Iersel, M. Jones, V. Moulton, and L. Veenema Nipius, When are quarnets sufficient to reconstruct semi-directed phylogenetic networks?, Bulletin of Mathematical Biology, 87 (2025), p. 136, 10.1007/s11538-025-01510-5.

[23] A. K. Kuchibhotla, J. E. Kolassa, and T. A. Kuffner, Post-selection inference, Annual Review of Statistics and Its Application, 9 (2022), pp. 505–527, 10.1146/annurev-statistics-100421-044639.

[24] B. Legried, E. K. Molloy, T. Warnow, and S. Roch, Polynomial-time statistical estimation of species trees under gene duplication and loss, Journal of Computational Biology, 28 (2021), pp. 452–468, 10.1089/cmb.2020.0424.

[25] J. Lescroart, A. Bonilla-Sánchez, C. Napolitano, D. L. Buitrago-Torres, H. E. Ramírez-Chaves, P. Pulido-Santacruz, W. J. Murphy, H. Svardal, and E. Eizirik, Extensive phylogenomic discordance and the complex evolutionary history of the neotropical cat genus Leopardus, Molecular Biology and Evolution, 40 (2023), 10.1093/molbev/msad255.

[26] Y. Liu and J. Xie, Cauchy combination test: A powerful test with analytic p-value calculation under arbitrary dependency structures, Journal of the American Statistical Association, 115 (2020), pp. 393–402, 10.1080/01621459.2018.1554485. PMID: 33012899.

[27] A. Markin and O. Eulenstein, Quartet-based inference is statistically consistent under the unified duplication-loss-coalescence model, Bioinformatics, 37 (2021), pp. 4064–4074, 10.1093/bioinformatics/btab414.

[28] S. Mirarab, Species tree estimation using ASTRAL: Practical considerations, in Species tree inference: A Guide to Methods and Applications, L. Kubatko and L. L. Knowles, eds., Princeton University Press, Princeton, 2023, ch. 3, pp. 43–67, 10.1515/9780691245157-007.

[29] S. Mirarab, R. Reaz, M. S. Bayzid, T. Zimmermann, M. S. Swenson, and T. Warnow, ASTRAL: genome-scale coalescent-based species tree estimation, Bioinformatics, 30 (2014), pp. i541–i548, 10.1093/bioinformatics/btu462.

[30] J. D. Mitchell, E. S. Allman, and J. A. Rhodes, Hypothesis testing near singularities and boundaries, Electronic Journal of Statistics, 13 (2019), pp. 2150–2193, 10.1214/19-EJS1576.

[31] A. Rafi, A. M. S. Rumi, S. A. Hakim, Sohaib, M. T. Tahmid, R. J. I. Momin, T. A. Zaman, R. Reaz, and M. S. Bayzid, wQFM-TREE: highly accurate and scalable quartet-based species tree inference from gene trees, Bioinformatics Advances, 5 (2025), p. vbaf053, 10.1093/bioadv/vbaf053.

[32] J. A. Rhodes, H. Baños, J. D. Mitchell, and E. S. Allman, MSCquartets 1.0: quartet methods for species trees and networks under the multispecies coalescent model in R, Bioinformatics, 37 (2020), pp. 1766–1768, 10.1093/bioinformatics/btaa868.

[33] J. A. Rhodes, H. Baños, J. Xu, and C. Ané, Identifying circular orders for blobs in phylogenetic networks, Advances in Applied Mathematics, 163 (2025), p. 102804, 10.1016/j.aam.2024.102804.

[34] G. Schwarz, Estimating the dimension of a model, The Annals of Statistics, 6 (1978), pp. 461–464.

[35] J. Shao, Mathematical Statistics, Springer Texts in Statistics, Springer, New York, NY, 2nd ed., 2003, 10.1007/b97553.

[36] C. Solís-Lemus and C. Ané, Inferring Phylogenetic Networks with Maximum Pseudolikelihood under Incomplete Lineage Sorting, PLoS Genetics, 12 (2016), p. e1005896, 10.1371/journal.pgen.1005896.

[37] C. Solís-Lemus, M. Yang, and C. Ané, Inconsistency of species tree methods under gene flow, Systematic biology, 65 (2016), pp. 843–851, 10.1093/sysbio/syw030.

[38] J. Taylor and R. Tibshirani, Post-selection inference for ℓ1-penalized likelihood models, Canadian Journal of Statistics, 46 (2018), pp. 41–61, 10.1002/cjs.11313.

[39] J. Taylor and R. J. Tibshirani, Statistical learning and selective inference, Proceedings of the National Academy of Sciences, 112 (2015), pp. 7629–7634, 10.1073/pnas.1507583112.

[40] L. H. C. Tippett, The Methods of Statistics. An introduction mainly for workers in the biological sciences., Williams and Norgate, Ltd., London., 1931.

[41] J. Xu and C. Ané, Identifiability of local and global features of phylogenetic networks from average distances, Journal of Mathematical Biology, 86 (2023), p. 12, 10.1007/s00285-022-01847-8.

[42] C. Zhang and S. Mirarab, Weighting by gene tree uncertainty improves accuracy of quartet-based species trees, Molecular Biology and Evolution, 39 (2022), p. msac215, 10.1093/molbev/msac215.

[43] C. Zhang, M. Rabiee, E. Sayyari, and S. Mirarab, ASTRAL-III: polynomial time species tree reconstruction from partially resolved gene trees, BMC Bioinformatics, 19 (2018), p. 153, 10.1186/s12859-018-2129-y.

