## Supplementary Materials for "Statistical inference of the Tree of Blobs of a phylogenetic network from quartet concordance factors"

Code to reproduce simulated data and ECToBlob analyses is at https://github.com/eallman/2026_ECToBlob_Sims.

Network figures were drawn with [3, 1].

### A. Simulations to assess the accuracy of ECToBlob.

**Model network.** Figure S1 shows the model network $N$ used for simulations (left), and its true tree of blob $T_B(N)$ (right). The network's branch lengths, in coalescent units drawn from $\Gamma(20, 1/20)$, were scaled by factor $s = 1$, 0.75, 0.5, 0.25 to give increasing levels of incomplete lineage sorting. Inheritance parameters $\gamma > 0.5$ are shown in the figure. `ECToBlob` was applied with the default values $\beta = 0.8$ and $\alpha = 0.05$.

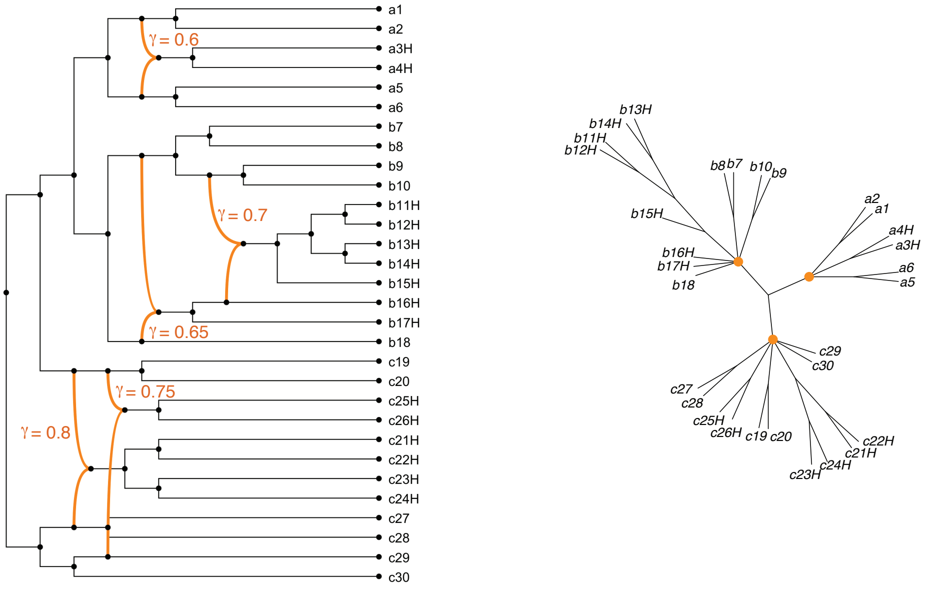


**Figure S1.** *The model network $N$ (left) used for simulations, and its tree of blobs $T_B(N)$ (right). $N$ is planar, but not outer-labelled planar, with three blobs: a 4-cycle leading to hybrid taxa $\{a3H, a4H\}$, a level-2 7-blob with one reticulation descendant of another, leading to hybrid taxa $\{b11H, \ldots, b17H\}$, and another level-2 7-blob leading to hybrid taxa $\{c21H, \ldots, c26H\}$. The blobs in $N$ correspond to nodes of degree 4, 7, and 7 in $T_B(N)$. See code repository for edge lengths on $N$.*

*University of Alaska - Fairbanks
†University of Wisconsin - Madison
‡California State University - San Bernardino

### Accuracy of `ECToBlob` in simulations.

**Table S1**

*Accuracy of `ECToBlob`, measured as the number of replicates (out of 100, from each number $m$ of gene trees and branch length scaling $s$) for which the true tree of blobs $T_B(N)$ was inferred correctly by `ECToBlob`. For each multiple test correction method (*Bon, BBC, Cauchy *and* CBC*), the value indicates the minimum accuracy over the three qType methods (`bi`, `mul`, and `quad`). Data were simulated under the network in Fig. S1, after scaling all branch lengths by a factor of $s$. A smaller scaling factor $s$ corresponds to increased incomplete lineage sorting and more gene tree discordance.*

| | 1000 gene trees | | | | 500 gene trees | | | |
|---|---|---|---|---|---|---|---|---|
| $s$ | Bon | BBC | Cauchy | CBC | Bon | BBC | Cauchy | CBC |
| 1.0 | 99 | 100 | 67 | 75 | 100 | 100 | 0 | 0 |
| 0.75 | 99 | 100 | 5 | 6 | 92 | 84 | 0 | 0 |
| 0.5 | 89 | 87 | 0 | 0 | 26 | 14 | 0 | 0 |
| 0.25 | 3 | 2 | 0 | 0 | 0 | 0 | 0 | 0 |

| | 300 gene trees | | | |
|---|---|---|---|---|
| $s$ | Bon | BBC | Cauchy | CBC |
| 1.0 | 86 | 81 | 0 | 0 |
| 0.75 | 42 | 37 | 0 | 0 |
| 0.5 | 1 | 1 | 0 | 0 |
| 0.25 | 0 | 0 | 0 | 0 |

#### Timing information for networks with 50, 100 taxa.

As a test, we ran ECToBlob on level-2 binary networks with 50 and 100 tips. The 50-taxon network $N_{50}$ has two blobs, and the 100-taxon network was formed by joining two copies of $N_{50}$ so had four blobs. The Newick strings for these networks are available on the GitHub site for Supplementary Materials. Timing information is given in the tables below.

**Table S2**

*Time (in secs) to compute the* `ECToBlob` *tree on a 50-taxon network and a single simulated 500 gene tree dataset. Analyses were performed with a Macbook Pro M3 chip with 36 Gb memory.*

Gene tree processing:
read 500 gene trees, perform 230300 hypothesis tests

| $s = 1.0$ | $s = 0.75$ | $s = 0.5$ | $s = 0.25$ | Mean |
|---|---|---|---|---|
| 124.87 | 130.48 | 135.68 | 140.03 | 132.76 |

ECToBlob: 50-taxon network, 500 gene trees

| | $s = 1.0$ | | | | $s = 0.75$ | | | |
|---|---|---|---|---|---|---|---|---|
| | Bon | BBC | Cauchy | CBC | Bon | BBC | Cauchy | CBC |
| `bi` | 49.54 | 49.67 | 50.12 | 47.70 | 53.97 | 52.56 | 51.14 | 50.46 |
| `mul` | 30.84 | 69.08 | 118.60 | 31.07 | 39.22 | 70.68 | 123.52 | 38.80 |
| `quad` | 16.67 | 35.09 | 65.79 | 16.67 | 17.99 | 32.06 | 59.91 | 17.86 |

ECToBlob: 50-taxon network, 500 gene trees

| | $s = 0.5$ | | | | $s = 0.25$ | | | |
|---|---|---|---|---|---|---|---|---|
| | Bon | BBC | Cauchy | CBC | Bon | BBC | Cauchy | CBC |
| `bi` | 53.59 | 51.20 | 48.27 | 46.62 | 59.46 | 53.48 | 46.36 | 47.97 |
| `mul` | 35.88 | 65.01 | 97.29 | 36.60 | 39.42 | 84.86 | 122.56 | 43.09 |
| `quad` | 20.77 | 31.91 | 55.32 | 20.76 | 24.68 | 36.17 | 51.26 | 23.29 |

**Table S3**

*Time (in hours) to compute the* `ECToBlob` *tree on a 100-taxon network and a single simulated 500 gene tree dataset. Analyses were performed with a Macbook Pro M3 chip with 36 Gb memory.*

Gene tree processing:
read 500 gene trees, perform 3921225 hypothesis tests

| | |
|---|---|
| $s = 1.0$ | 0.71 hours |

ECToBlob: 100-taxon network, 500 gene trees

| | $s = 1.0$ | | | |
|---|---|---|---|---|
| | Bon | BBC | Cauchy | CBC |
| `bi` | 2.46 | 2.46 | 2.48 | 2.43 |
| `mul` | 1.28 | 8.80 | 16.20 | 1.27 |
| `quad` | 0.20 | 0.42 | 0.69 | 0.20 |

#### B. Simulation for the comparison to TOB-QMC.

We ran a simulation experiment to compare `TOB-QMC` [2] and `ECToBlob`. The performance of these methods depends on several options, including the choice of parameters $\beta$, $\alpha$. However, while conceptually related, the tests for which these parameters are used as levels differ: the individual star and T3 test levels for `TOB-QMC`; the corrected star and corrected T1 for `ECToBlob`. Therefore, we first compare the two methods under a fixed common choice of these numerical parameter settings, and then at their optimal performance values across all choices.

For this, we repeated the following simulation procedure 100 times: We simulated 1000 gene trees using `PhyloCoalsimulations` on the network whose topology is depicted in Figure S2. For this network, all branches were 0.5 coalescent units, and all inheritance probabilities were set to 0.5.

We chose to investigate this network based on the following reasoning. The network has several blobs near the leaves, and none far from the leaves. Since edges near the leaves are associated to fewer quartets ($\mathcal{O}(n^2)$ or less) than those more centrally located ($\mathcal{O}(n^4)$ quartets), it is more likely some quartet tests for central edges of the initial resolution of the tree of blobs will, by chance, give low $p$-values even when those edges should not be contracted. The multiple test corrections of `ECToBlob` should address this, while `TOB-QMC` might be more prone to erroneous contractions.

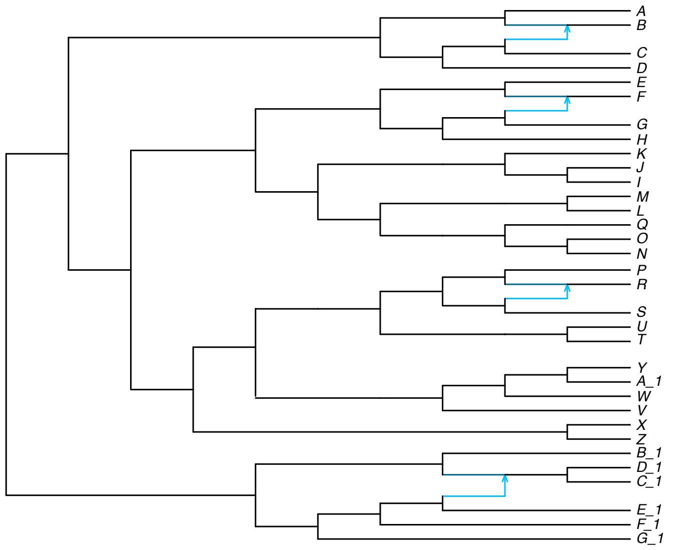


**Figure S2.** *Topology of the 33-taxon level-1 network used for the comparison between* `ECToBlob` *and* `TOB-QMC`*. All edge lengths were set to 0.5 coalescent units, and all inheritance probabilities were set to 0.5.*

Since both methods require a starting tree as input, we obtained a common one by running `ASTRAL-IV` [4] on each dataset. In all cases, we confirmed that this tree was a refinement of $T_B(N)$. We also set $\beta = 1$ (the test level for a star-like quartet) for each method. This ensures that no quartet is judged as polytomous, and thus allows us to focus on and compare the effect of edge contraction due to blobs.

For the first experiment, we ran both methods on each of the 100 datasets using a fixed

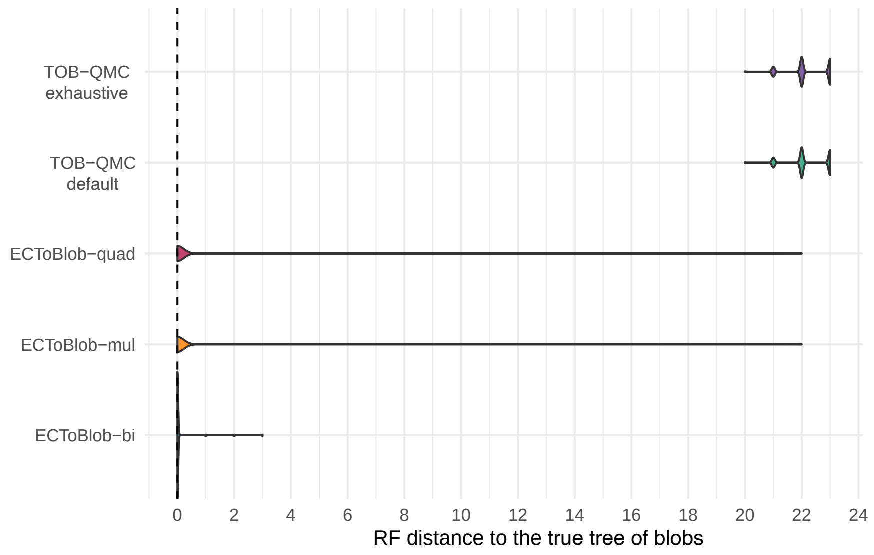


**Figure S3.** *Violin plots showing the RF distances to the true tree of blobs, with $\alpha = 0.05$ for each method. For `ECToBlob`, we used the* Bon *test correction across all three quartet-type choices. 'TOB-QMC default' denotes runs in which the search for the minimum p-value was performed using default settings, whereas 'TOB-QMC exhaustive' denotes runs in which this search was exhaustive. Each violin summarizes 100 replicates.*

value of $\alpha = 0.05$. We ran `TOB-QMC` twice, first using the setting `--iter_limit_blob 0` to perform an exhaustive search for the minimum $p$-value, and second, using the default setting for the number of iterations. We ran `ECToBLob` using Bon for multiple-test correction, each quartet type (`bi`, `mul`, `quad`), and the `early` selection rule.

Figure S3 displays the RF distance to the true tree of blobs for `TOB-QMC` and `ECToBlob` across each quartet-type choice. This plot shows that, at significance level $\alpha = 0.05$, `TOB-QMC` recovers a tree far from the true tree of blobs, whereas `ECToBlob` is able to recover the true one many times under all choices of quartet type.

Note this should not be interpreted as showing `TOB-QMC` necessarily performs poorly. Rather, it suggests that `TOB-QMC` may require a more extreme value of $\alpha$ than empiricists might choose based on general familiarity with hypothesis testing levels, likely due to the absence of multiple-testing correction. It also motivates our second experiment.

We next investigate the optimal performance of both methods across all choices of $\alpha$, using the same 100 datasets.

For `TOB-QMC`, each replicate dataset was analyzed as follows. 1) Following the developers' guidelines, we annotated all branches of the starting tree with the minimum $p$-value found from T3 hypothesis testing of single quartets, by running `TOB-QMC` using an exhaustive search and the default setting for $\alpha$. 2) A user might then choose another fixed test level $\alpha$ and have `TOB-QMC` contract all edges whose minimum $p$-values are less than $\alpha$. We instead explored all possible output trees by ordering the minimum edge $p$-values and making choices of $\alpha$ between consecutive elements. 3) We computed the Robinson-Foulds (RF) distance between each of the output trees and the true tree of blobs, recording the minimum RF distance as a measure

**Table S4**

*Distribution of the RF distance between the true tree of blobs and the best estimate (over all $\alpha$ values for each replicate). For each method, the values give the number of replicates at this RF distance across a total of 100 replicates. For `ECToBlob`, we used the* Bon *test correction across all three quartet-type choices.*

| RF distance | ECToBlob bi | ECToBlob mul | ECToBlob quad | TOB-QMC default | TOB-QMC exhaustive |
|---|---|---|---|---|---|
| 0 | 100 | 100 | 100 | 55 | 55 |
| 1 | 0 | 0 | 0 | 45 | 45 |

of the method's optimal accuracy.

For `ECToBLob`, we followed a similar procedure. 1) From the starting tree and for each quartet type (`bi`, `mul`, `quad`) we ran `ECToBLob` using Bon for multiple-test correction. The output includes a sequence of trees obtained after contracting edges of minimal $p$-value. Typically, a user would choose a fixed test level $\alpha$ and either the `early` or `late` selection rule to select the estimated tree in the sequence (as done in the previous experiment). Here we used the `early` selection rule and explored all estimates that could be obtained from all choices of $\alpha$ by ordering the $p_k$ of the trees $T_k$ in the output sequence of `ECToBLob`, and choosing a set of $\alpha$ values between consecutive elements. 2) For each quartet type, we computed the RF distance between each tree obtained in this way and the true tree of blobs, recording the minimum distance.

Table S4 shows the RF distance to the true tree of blobs obtained from `TOB-QMC` and `ECToBlob`, which was either 0 or 1 in all cases. This underscores our interpretation of the first experiment, indicating $\alpha = 0.05$ may be far from an optimal choice for `TOB-QMC`. `ECToBlob` always recovered the true tree of blobs $T_B(N)$, whereas `TOB-QMC` recovered $T_B(N)$ only about half of the time.

We suspect that the `ECToBlob`'s improved performance over `TOB-QMC` in this experiment is primarily due to its correction of $p$-values for multiple testing. However, using the T1 rather than the T3 test might also be a factor, as could `ECToBlob`'s computation of a single $p$-value from retained cut edges to judge its output tree.

### REFERENCES


[1] C. Ané, *PhyloPlots*, 2025, https://github.com/JuliaPhylo/PhyloPlots.jl. v. 2.1.0.

[2] J. Dai, Y. Han, and E. K. Molloy, *Quartet-based species tree methods enable fast and consistent tree of blobs reconstruction under the network multispecies coalescent*, bioRxiv, version 5 (2026), https://doi.org/10.1101/2025.11.05.686850.

[3] D. H. Huson, *Sketch, capture and layout phylogenies*, PLOS Computational Biology, 21 (2025), https://doi.org/10.1371/journal.pcbi.1013805. v. 2.2.11.

[4] C. Zhang, R. Nielsen, and S. Mirarab, *ASTER: A package for large-scale phylogenomic reconstructions*, Molecular Biology and Evolution, 42 (2025), p. msaf172, https://doi.org/10.1093/molbev/msaf172.
